# Androgens at the skin surface regulate *S. aureus* pathogenesis through the activation of *agr* quorum sensing

**DOI:** 10.1101/2024.02.10.579753

**Authors:** Maria Sindhura John, Mahendran Chinnappan, Methinee Artami, Mohini Bhattacharya, Rebecca A. Keogh, Jeffrey Kavanaugh, Tripti Sharma, Alexander R. Horswill, Tamia A. Harris-Tryon

## Abstract

*Staphylococcus aureus,* the most frequent cause of skin infections, is more common in men than women and selectively colonizes the skin during inflammation. Yet, the specific cues that drive infection in these settings remain unclear. Here we show that the host androgens testosterone and dihydrotestosterone promote *S. aureus* pathogenesis and skin infection. Without the secretion of these hormones, skin infection *in vivo* is limited. Testosterone activates *S. aureus* virulence in a concentration dependent manner through stimulation of the *agr* quorum sensing system, with the capacity to circumvent other inhibitory signals in the environment. Taken together, our work defines a previously uncharacterized inter-kingdom signal between the skin and the opportunistic pathogen *S. aureus* and identifies the mechanism of sex-dependent differences in *S. aureus* skin infection.

**One-Sentence Summary:** Testosterone promotes *S. aureus* pathogenesis through activation of the *agr* quorum sensing system.

## Introduction

The community of commensal organisms residing in mammalian skin thrive in the desiccate, acidic, and lipid-rich landscape (*1*). Staphylococcal species such as *S. epidermidis* and *S. hominis* are established resident microbes of human skin (*2, 3*). In contrast, the related opportunistic pathogen *S. aureus* is restricted to the nares of only a third of humans, with skin colonization and infection occurring as a rare event (*4*). Indeed, the numerous virulence factors encoded by *S. aureus* that allow it to bind, invade, and damage epithelial cells and cause infection are dampened by coordinated regulatory networks that tune their expression depending on sensed environmental conditions (*5*).

The most well-described system of virulence regulation in *S. aureus* is the accessory gene regulator (*agr)* quorum-sensing system (*5, 6*) that controls the expression of toxins and adhesins required for skin colonization. *S. aureus* strains can auto-activate *agr* signaling through the production of an autoinducing peptide (AIP) signal. They have also evolved the ability to respond to quorum signals generated by other bacteria, including the variant AIP signals from non-cognate strains of *S. aureus* (*5*). Though similar in structure, many non-cognate AIPs inhibit quorum sensing in *S. aureus* (*5*). *Agr* quorum sensing can also be inhibited by multiple host derived mechanisms (*7–9*).

It remains unclear what host cues promote *S. aureus* pathogenesis and allow it to begin to effectively colonize the skin during flares of inflammatory skin conditions, such as atopic dermatitis, or why males are more susceptible to skin and soft tissue infections with *S. aureus* than females (*10–15*). One established difference between males and females is the amount of androgens generated by the skin and by sex organs (*16, 17*). It was recently reported that IL-4 mediated inflammation in atopic dermatitis regulates androgen production in the skin, providing a potential link between the sexual dimorphism of *S. aureus* infections and the blooms of *S. aureus* colonization during IL-4 mediated skin conditions (*18, 19*).

Here, we demonstrate that mice that secrete lower levels of testosterone and dihydrotestosterone (DHT) are resistant to *S. aureus* skin infection. Further, testosterone and DHT specifically activate the *agr* quorum sensing system, independently of the bacterially derived agonist AIP-I. Testosterone can overcome inhibitory signals from other bacterial strains in a dose dependent manner, indicating that testosterone may be a host-derived cue that directly antagonizes inhibitory signals from other competing bacteria at the skin surface. Our findings are further supported by protein folding models that demonstrate a testosterone binding pocket at a separate site from AIP-I on the *agr* receptor, AgrC. Taken together, our work defines a previously uncharacterized interaction between the skin and the opportunistic pathogen, *S. aureus,* and defines a molecular inter-kingdom signal between the skin and the microbiota.

## Results

### Androgen secretion at the skin surface is required for *S. aureus* skin infection

Higher androgen secretion is associated with *S. aureus* colonization and infection (*4, 13, 20*). *S. aureus* colonization is significantly more common in men versus women (*10–13*) and *S. aureus* induces greater necrosis in male murine skin infections compared to female mice (*13*). To confirm these findings in a model of methicillin resistant *S. aureus* (MRSA) skin infection, we epicutaneously infected male and female age-matched C57BL6 mice with a bioluminescent strain of MRSA (MRSA::*lux*), which generates light in proportion to the number of colony forming units present at the skin surface (*21*). Consistent with prior findings, we observed a two log fold difference in MRSA infection in male mice compared to female mice (Fig. 1A and fig. S1) (*13*). In keeping with our prior study in humans (*17*), male mice also secrete greater amounts of androgens at the skin surface compared to female mice (Fig. 1B, C). Taken together, these findings demonstrate the association between higher androgen secretion and increased bacterial burdens during *S. aureus* skin infection.

**Fig. 1.**
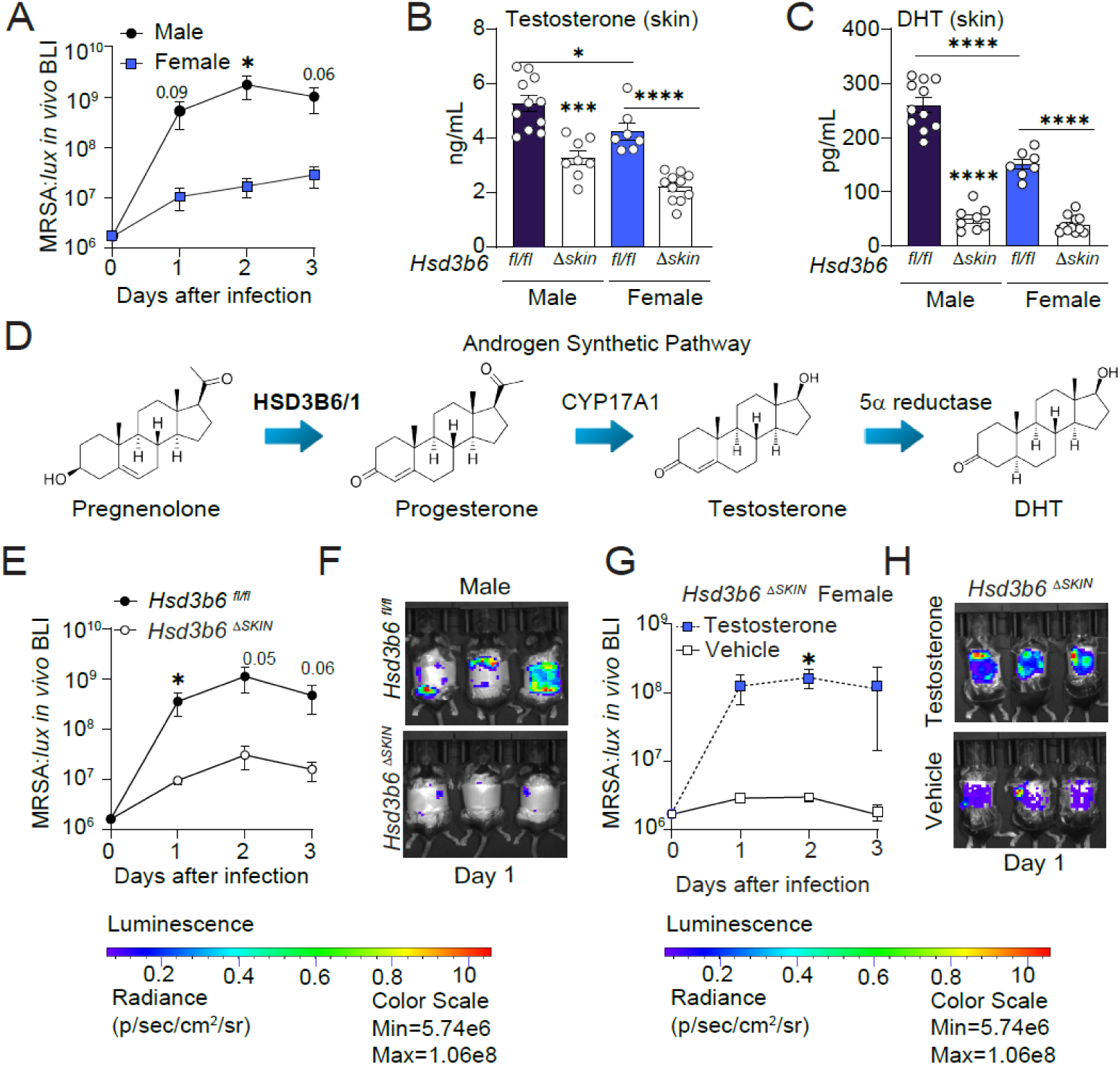
*Hsd3b6^Δskin^* mice lack skin-secreted androgens and are resistant to skin infection with *S. aureus.* **(A)** Male and female wild-type (WT) mice were epicutaneously infected for 3 days with 1×10^6^ colony forming units (CFUs) of a bioluminescent strain of methicillin resistant *S. aureus* (MRSA::*lux*). Bioluminescence quantified over time. *n*=6 male and *n=*7 female mice, aggregate of two experiments. **(B, C)** Testosterone (B) and dihydrotestosterone (DHT) (C) quantification from the skin secretions of male *Hsd3b6^fl/fl^* (*n*=11) and *Hsd3b6^Δskin^* (*n*=8) and female *Hsd3b6^fl/fl^*(*n*=7) and *Hsd3b6^Δskin^* (*n*=11) mice, age-matched at 7 weeks. **(D)** Schematic of HSD3B6/1 enzyme mediated conversion of pregnenolone to testosterone and DHT. **(E)** Male *Hsd3b6^fl/fl^*(*n*=5) and *Hsd3b6^Δskin^* (*n*=7) mice epicutaneously infected for 3 days with 1×10^6^ CFUs of MRSA::*lux*. Aggregate of two experiments **(F)** Representative image of (E). **(G)** Female *Hsd3b6^Δskin^* mice epicutaneously infected for 3 days with 1×10^6^ CFUs of MRSA:*:lux* treated with testosterone or vehicle control. *n=5* per group. Aggregate of two experiments **(H)** Representative image of (G). Means ± SEM (error bars) are plotted. \**p* < 0.05; \*\**p* < 0.01; \*\*\**p* < 0.001, \*\*\*\**p* <0.0001, ns, not significant by two-tailed *t*-test.

In addition to the increased burden of *S. aureus* in male patients, people with the inflammatory skin condition atopic dermatitis are ubiquitously colonized by *S. aureus* (*22–25*), with little understanding as to why *S. aureus* begins to dominate the skin surface in this setting (*22–25*). Interestingly, the immune system in atopic dermatitis can directly regulate androgen production through stimulation of a rate limiting enzyme in the synthesis of steroid hormones, 3β-hydroxysteroid dehydrogenase 1 (HSD3B1) (*18, 26, 27*) (Fig. 1D). Thus, immune regulation of

HSD3B1 in atopic dermatitis provides a second link between high androgen states and *S. aureus* colonization (*18*). To further dissect the link between androgens and *S. aureus in vivo* we created mice lacking androgen production at the skin surface, through skin specific deletion of the mouse ortholog to *HSD3B1, Hsd3b6* (*18*). We used CRISPR/Cas-9 mediated gene targeting to insert *loxP* sites around the first exon of the *Hsd3b6* locus creating the *Hsd3b6^fl/fl^* mice (fig. S2A). We then crossed these mice to a skin specific Cre driver (*K14-Cre^+/-^*) to generate the *Hsd3b6^Δskin^ (K14Cre^+-/^; Hsd3b6^fl/fl^)* mice and verified the loss of HSD3B6 expression by immunofluorescence (fig. S2B). *Hsd3b6^Δskin^* mice displayed no visible phenotypes when reared in a specified-pathogen-free (SPF) facility and displayed no signs of skin inflammation (fig. S2C). However, *Hsd3b6^Δskin^* mice had marked reductions in the amount of testosterone, progesterone, and DHT secreted at the skin surface compared to *Hsd3b6^fl/fl^* mice (Fig. 1B-C, fig. S3A, B). *Hsd3b6^Δskin^* mice did not display differences in serum production of hormones, weights, immune cell populations, or skin barrier function (fig. S3C-F).

Next, we assessed the susceptibility of *Hsd3b6^Δskin^* mice to skin infection. Epicutaneous infection of the *Hsd3b6^Δskin^* mice with MRSA::*lux* resulted in a marked reduction of MRSA skin infection compared to the *Hsd3b6^fl/fl^* control mice infected with the same inoculum (Fig. 1E-F, fig. S4A). Additionally, infection of female *Hsd3b6^Δskin^* mice with MRSA::*lux* was augmented by the topical addition of testosterone at the skin surface (Fig. 1G-H, fig. S4B). Thus, the reduction of testosterone, DHT, and progesterone at the skin surface protected the skin from skin infection and treatment with exogenous testosterone promoted *S. aureus* infection of the skin. Reduction of skin secreted hormones also abrogated sex-dependent differences in infection (Fig. 1E-H).

### Testosterone and DHT activate *agr* quorum sensing and promote *S. aureus* pathogenesis

To determine how hormones might regulate the *S. aureus* transcriptome, we next sequenced RNA from *S. aureus* treated with testosterone compared to controls. Interestingly, testosterone had a very narrow impact on the *S. aureus* regulon, with marked increases in the expression of a few genes, *agrB, agrD, agrC, agrA, psmα, psmβ,* and *RNAIII* (Fig. 2A), all of which are in the accessory gene regulator (*agr)* quorum-sensing pathway (*5, 6, 28, 29*) (Fig. 2B). In contrast, pregnenolone, a hormone with a similar structure and carbon count to testosterone, had no discernable impact on the *S. aureus* transcriptome (fig. S5A, B).

**Fig. 2.**
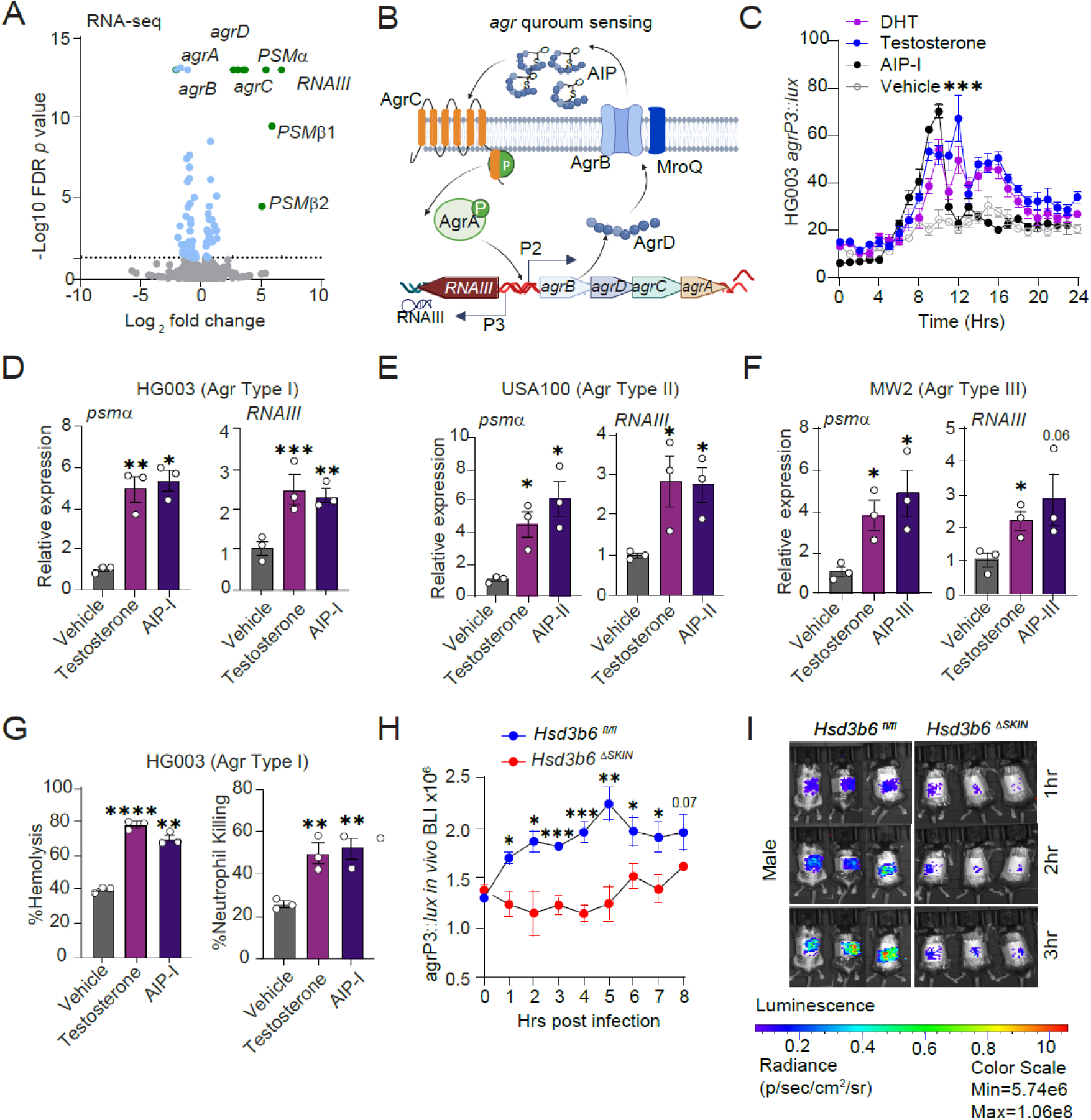
Testosterone stimulates *agr* quorum sensing in *S. aureus.* **(A)** HG003 strain of *S. aureus* was treated with 10nM testosterone or untreated with vehicle alone. Volcano plot demonstrating genes with > 4-fold change in expression (green) after treatment as determined by transcriptomics (RNA-seq). **(B)** Schematic of *S. aureus agr* two-component quorum sensing system. **(C)** Bioluminescence of *agr*-P3 reporter (HG003*agrP3::lux*) treated with testosterone, DHT, AIP-I, or vehicle alone. Statistics of testosterone compared to vehicle at 12 hours. **(D)** qRT-PCR of *S. aureus* strain, HG003 (Type I strain), cultured to mid-exponential growth treated with 10nM Testosterone, AIP-I, or untreated with vehicle alone (*n*=3). Gene expression of target gene normalized to *gyrA* expression. **(E, F)** qRT-PCR of *agr* Type II (USA100) (E) and Type III (MW2) (F) *S. aureus* strains cultured to mid-exponential growth treated with 10 nM Testosterone, AIP-II and AIP-III respectively (*n*=3). Gene expression of target gene normalized to *gyrA* expression. **(G)** Percentage of *S. aureus* induced red blood cell (RBC) hemolysis and neutrophil killing with and without 10nM Testosterone or AIP-I. **(H, I)** In vivo analysis of *agr-*P3 reporter. Epicutaneous infection of male Hsd3b6*^fl/fl^*(*n*=7) and Hsd3b6^Δskin^ mice (*n*=6) with HG003*agrP3::lux* with bioluminescence quantified over time. Aggregate of two experiments (H) and representative images (I). Means ± SEM (error bars) are plotted.\**p* < 0.05; \*\**p* < 0.01; \*\*\**p* < 0.001, \*\*\*\**p* <0.0001, ns, not significant by two-tailed unpaired *t*-test.

*Agr* activation occurs through transcriptional regulation of the P3 promoter (Fig. 2B) (*5*). We therefore tested the impact of androgens on luminescent P3 promoter fusions of *S. aureus* (HG003 *agrP3*::*lux*) that generate bioluminescence in proportion to the activation of quorum sensing (*30*). In this model, testosterone and DHT activated the *agr* P3 promoter, with similar kinetics to the established ligand AIP-I (Fig. 2C). In contrast, estradiol and progesterone had no impact on *agr* activation (fig. S5C-F). To confirm these findings, we quantified the transcription of key readouts of *agr* activity, *psmα* and *RNAIII,* in the HG003 strain of *S. aureus* and demonstrated that testosterone stimulates the expression of both transcripts (Fig. 2D). Thus, testosterone and DHT activate the transcription of the *agr* regulon in *S. aureus*, but estradiol, pregnenolone, and progesterone do not.

Every staphylococcal isolate contains only a single copy of the *agr* system, and each species produces different types of autoinducing AIP signal through variation in the *agrBDCA* operon (*5*). There are four types of AIP signal made by *S. aureus* and HG003 falls into the *agr* Type I class. To test the generality of the effects of testosterone across *agr* types, we treated additional strains with testosterone and measured *psm*α and *RNAIII* expression, including USA100 (Type II) and MW2 (Type III) (*31–34*). All strains showed robust expression of *psm*α and *RNAIII* (Fig. 2E, F). Testosterone also activated the P3 promoter in Type II and Type III luminescent strains (fig. S5G, H). Further, testosterone stimulated the transcription of the *agr* regulated virulence factors *luk*S*-PV*, *hla*, *hld*, the cytoplasmic regulator *agrA* (fig. S6A-D) and increased red blood cell hemolysis and neutrophil killing capacity of *S. aureus* (Fig. 2G). These effects were comparable to those of the established *agr* ligand AIP-I (Fig. 2C-G, fig. 6A-D). Greater than 90% of strains are Type I-III (*35, 36*), suggesting that testosterone stimulates virulence across *S. aureus* strains with active *agr* systems. Given the strong association between *S. aureus* and atopic dermatitis (*37*), we also tested an array of strains obtained from diseased skin (*25*). Testosterone treatment increased the transcription of *psmα, RNAIII,* and *agrA* in strains obtained from atopic dermatitis skin (fig. S6E-G).

Consistent with our prior *in vivo* data (Fig. 1E), quorum sensing was quenched in *Hsd3b6^Δskin^* mice infected with the quorum sensing reporter strain in comparison to *Hsd3b6^fl/fl^* mice (Fig. 2H, I. fig. S7A, B). *Agr* activation was also greater in male mice compared to female mice (fig. S7C). Taken together, these data show that the androgens testosterone and DHT stimulate the *agr* quorum sensing system and promote *S. aureus* infection *in vivo*.

### Androgens stimulate *agr* independently of the auto-inducing peptides

AIP, the established endogenous ligand of *agr*, is synthesized by the coordinated action of integral membrane endopeptidases AgrB and MroQ on the ribosomally generated propetide AgrD (Fig. 2B) (*28, 29, 38*). Thus, the biosynthesis mutant strain of *S. aureus, ΔagrBD*, lacks the ability to auto-stimulate the *agr* quorum sensing system (*9*). We hypothesized that testosterone would require AIP-I to activate *agr*. However, in the biosynthesis mutant, testosterone retained the ability to activate *agr* associated phenotypes, including stimulation of P3 promoter transcripts, increased hemolysis and neutrophil killing (Fig. 3A-C). These effects were dose dependent, with greater concentrations of testosterone increasing transcription of *RNAIII, agrA, agrC*, and *psmα* (fig. S8A). Additionally, we generated a biosynthesis mutant luminescent reporter (*ΔagrBD::lux)* and confirm that progesterone and estradiol had no effect on *S. aureus.* DHT and testosterone both retained the capacity to stimulate bioluminescence (fig. S8B). Since AIP and testosterone could act independently, we next wanted to test the impact of both AIP-I and testosterone on *agr* signaling. Indeed treatment of *S. aureus* with testosterone had the capacity to augment AIP-I signaling in a dose-dependent manner (Fig. 3D, fig. S8C), establishing that testosterone may synergize with AIP signals to regulate *S. aureus* pathogenesis.

**Fig. 3.**
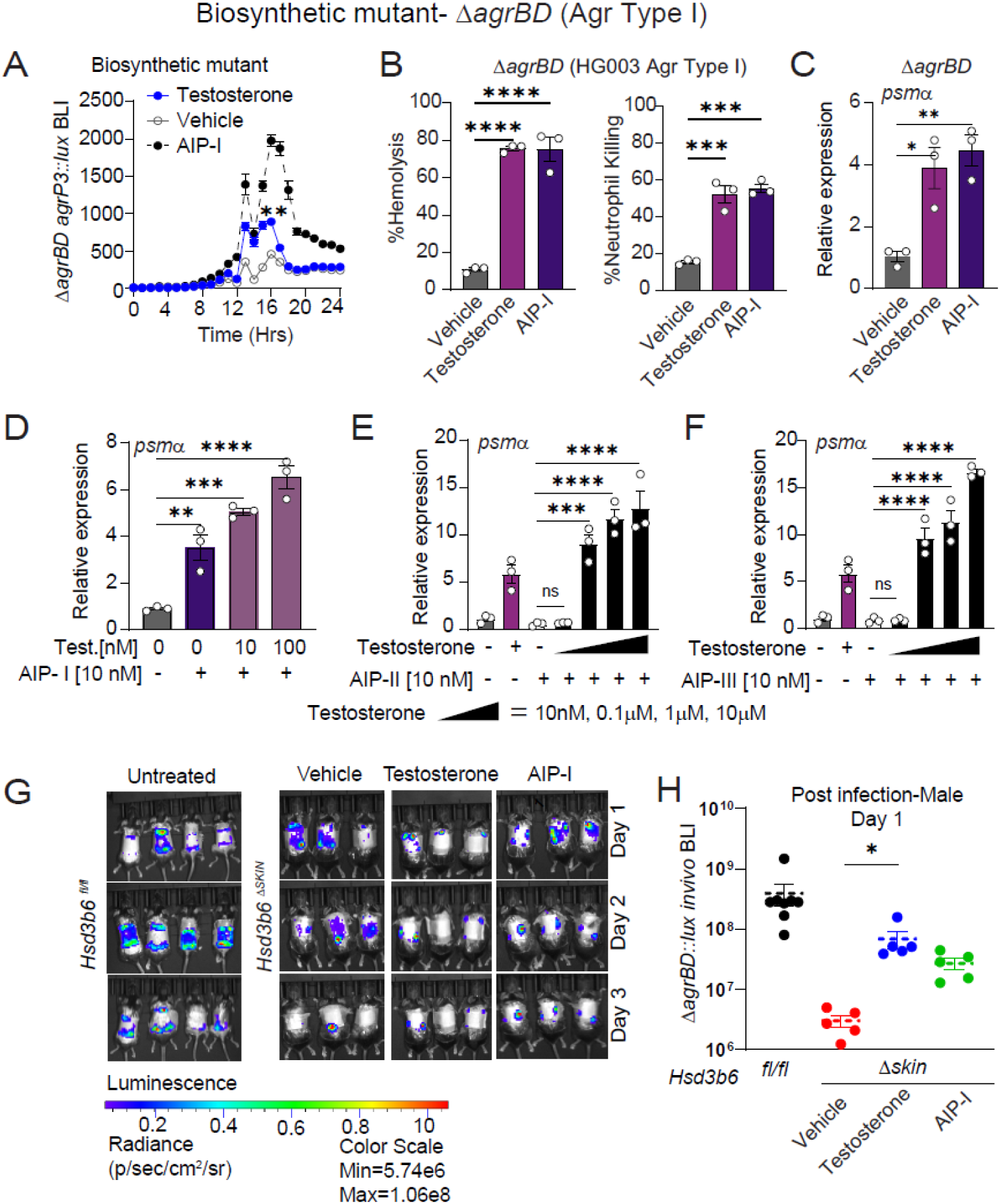
Testosterone retains *agr* activity in a biosynthetic mutant strain of *S. aureus*, *ΔagrBD*. **(A-C)** *S. aureus* biosynthetic mutant (*ΔagrBD-*HG003) and its respective *agr-P3* bioluminescent reporter strain (*ΔagrBD agrP3::lux*) were treated with 10nM Testosterone, AIP-I, or untreated. **(A)** *Agr-P3-lux* reporter kinetics were recorded every hour by a plate reader (*n*=12). Statistics of testosterone compared to vehicle. **(B)** Percentage of *S. aureus* induced hemolysis and neutrophil killing (*n*=3). **(C)** qRT-PCR of *psmα* expression normalized to *gyrA* expression. **(D)** qRT-PCR for *psmα* expression in the biosynthetic mutant strain treated with AIP-I and 10nM or 100nM or testosterone (*n*=3). **(E, F)** qRT-PCR for *psmα* expression in the biosynthetic mutant strain treated with AIP-II or AIP-III alone or in combination with increasing concentrations of testosterone (10nM to 10µM). **(G, H)** Male *Hsd3b6^fl/fl^* (*n*=8) and *Hsd3b6^Δskin^* mice were epicuataneously infected with 1×10^6^ CFUs of the biosynthetic mutant strain reporter (*ΔagrBD::lux*) treated topically with testosterone, AIP-I, or untreated (*n*=5) and bioluminescence quantified over time. Data from two experiments. Means ± SEM (error bars) are plotted.\**p* < 0.05; \*\**p* < 0.01; \*\*\**p* < 0.001, \*\*\*\**p* <0.0001, ns, not significant by two-tailed unpaired *t*-test.

Though cognate AIPs stimulate *agr,* non-cognate AIPs generated from other *agr* Types of *S. aureus* can inhibit *agr* signaling and are in development as *S. aureus* therapeutics (*5, 39*) (fig. S8D). Therefore, we tested how the stimulatory testosterone signal derived from the host might compete with inhibitory signals derived from competing *S. aureus* species that generate non-cognate AIPs, including AIP-II and AIP-III. Interestingly, when we exposed a Type-I strain to equal low nanomolar concentrations of AIP-II and testosterone, quorum sensing was inhibited, demonstrating that testosterone was unable to overcome inhibitory signals at the same concentration (Fig. 3E, fig. S9A). However, at higher concentrations, testosterone stimulated *agr* signaling and overcame the inhibitory AIP-II signal (Fig. 3E, fig. S9A). Similar dynamics were observed with AIP-III (Fig. 3F, fig. S9B). Taken together, these findings suggest that the host derived signal, testosterone, participates in the established crosstalk between competing microbes at the skin surface, and when present at levels higher than the inhibitor can overcome inhibitory signals generated towards *S. aureus*.

Lastly, we tested the effect of testosterone on the AIP biosynthesis mutant (*ΔagrBD::lux*) *in vivo*. Following epicutaneous infection, the biosynthesis mutant displayed a blunted infectious phenotype in the *Hsd3b6^Δskin^* mice compared to *Hsd3b6^fl/fl^* control (Fig. 3G, H). As was true *in vitro,* treatment with AIP-I or testosterone was able to boost *S. aureus* infection *in vivo* (Fig. 3G, H). These findings confirm that skin derived androgens facilitate epicutaneous infection in *S. aureus.* Moreover, both exogenous AIP and testosterone are sufficient to increase infection at the skin surface through enhanced expression of quorum sensing regulated virulence factors.

### The AgrC histidine kinase is required for testosterone mediated stimulation of *agr*

AIP stimulates the *agr* system through activation of the *AgrC* histidine kinase(*5, 40*). Once activated, AgrC phosphorylates the response regulator AgrA, which in turn autoinduces transcription of the Agr machinery (*5, 41*) (Fig. 2B). Given the specific impact of testosterone on the *agr* quorum sensing system (Fig. 2A) and its ability to cooperate and compete with established ligands of AgrC (Fig. 3)(*40*), we next tested if testosterone would require a complete AgrCA two-component system to regulate *S. aureus* virulence. We generated a constitutive bioluminescent reporter deficient in AgrC (*ΔagrC::lux)*(*9*). In contrast to the *ΔagrBD::lux, ΔagrC::lux* did not respond to testosterone (Figure 4A). Indeed, the inducing activity of testosterone on *S. aureus* required both AgrC and AgrA (Fig. 4B-D, fig. S10 A-C) and skin infections with the AgrC mutant had a muted infectious course (Fig. 4E,F, fig. S10D, E). Additionally, both *in vitro* and *in vivo* the addition of exogenous testosterone or AIP-I was not able to rescue *agr* activation in the *agrCA* deficient strains (Fig. 4B-D, fig. S10 A-C, F, G). Moreover, there were limited sex-differences in infections with *ΔagrC* (Fig. 4E, fig. S10E).

**Fig. 4.**
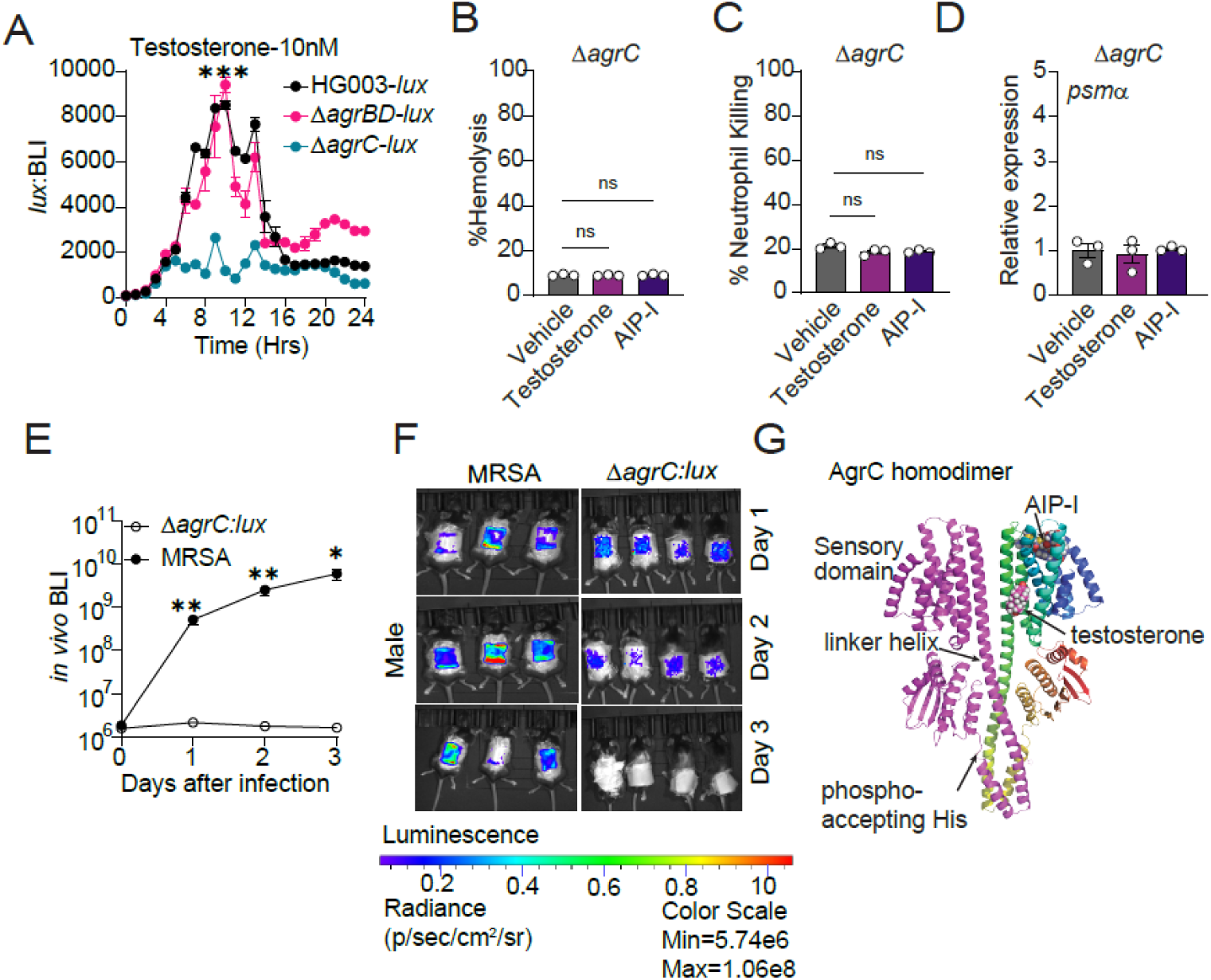
Testosterone does not stimulate quorum sensing in an AgrC mutant strain of *S. aureus*. **(A)** *S. aureus* wildtype (HG003*::lux*) and mutants (*ΔagrBD::lux and ΔagrC::lux)* strains treated with 10nM Testosterone. *lux* reporter bioluminescence kinetics recorded every hour by plate reader. Statistics of HG003 compared to *ΔagrC* treated with testosterone at 9 hours. **(B-D)** *ΔagrC* mutant treated with 10nM Testosterone, AIP-I, or untreated. Percentage of *S. aureus* induced hemolysis (B), neutrophil killing (C), and relative *psmα* expression (D) (*n*=3). **(E)** Male wild-type mice were epicutaneoulsy infected with 1×10^6^ CFUs of wild-type (*MRSA::lux*) (*n*=5) or agrC histidine kinase deficient (*ΔagrC::lux*) (*n*=6) bioluminescent reporter strain of *S. aureus.* Aggregate data of two experiments. **(F)** Representative bioluminescence images of (E). **(G)** Ribbon diagram of Alphafold2 structure of AgrC histidine kinase dimer with the docking of space filling models of testosterone and AIP-I and a representative testosterone docking solution. Means ± SEM (error bars) are plotted.\**p* < 0.05; \*\**p* < 0.01; \*\*\**p* < 0.001, \*\*\*\**p* <0.0001, ns, not significant by two-tailed unpaired *t*-test.

To gain further insight on interactions between AgrC and testosterone, we predicted the structure of the AgrC type I dimer using Alphafold2 (*42, 43*), and docked testosterone and AIP-I *in silico* on the AgrC sensory domain (*44–46*) (Fig. 4G, fig. S10H). Testosterone is predicted to bind to a hydrophobic cleft distinct from the established AIP binding stie. An allosteric binding site for testosterone is consistent with our studies demonstrating cooperative interactions between AIP-I and testosterone in Type-I strains (Fig. 3D-F). Taken together with our prior findings, these results demonstrate that testosterone enhances *S. aureus* pathogenesis and requires both the skin secretion of androgens and a functioning AgrCA two-component system.

### The enantiomer of testosterone inhibits *S. aureus* pathogenesis

As a final step to understanding the specificity of the interaction of testosterone with the *agr* system we tested the impact of a stereoisomer of testosterone, enantiomer-testosterone (*ent*-T), on *S. aureus* pathogenesis. Like other unnatural enantiomers, *ent*-T has the same physiochemical properties as testosterone but with a mirror image orientation at all six chiral centers that leads to altered rotation of polarized light (Fig. 5A) (*47, 48*). Enantiomers have been observed to have unique receptor binding and signaling properties (*48, 49*). Unlike the other hormones we tested which displayed neutral effects on the *agr* system (fig. S5C-F), *ent*-T acted as a broad inhibitor of the *agr* quorum sensing system (Fig. 5B-H, fig. S11). *Ent*-T was able to decrease *S. aureus* dependent hemolysis and neutrophil killing in Type-1 *agr* strains (Fig. 5B, C). Further, *ent*-T had the capacity to decrease the expression of *agr* transcriptional readouts in Type-II, Type-III, and atopic dermatitis strains of *S. aureus* (Fig. 5D-F, fig. S11A). Moreover, topical application of *ent*-T to the skin of *S. aureus* infected mice turns off quorum sensing *in vivo* (Fig. 5G, H; fig. S11B). Thus, *ent*-T is a global inhibitor of *S. aureus* quorum sensing that works across *agr* types, similar to established bacterially derived regulators of *agr,* such as non-cognate AIPs (Fig. 5B, C) (*6*).

**Fig. 5.**
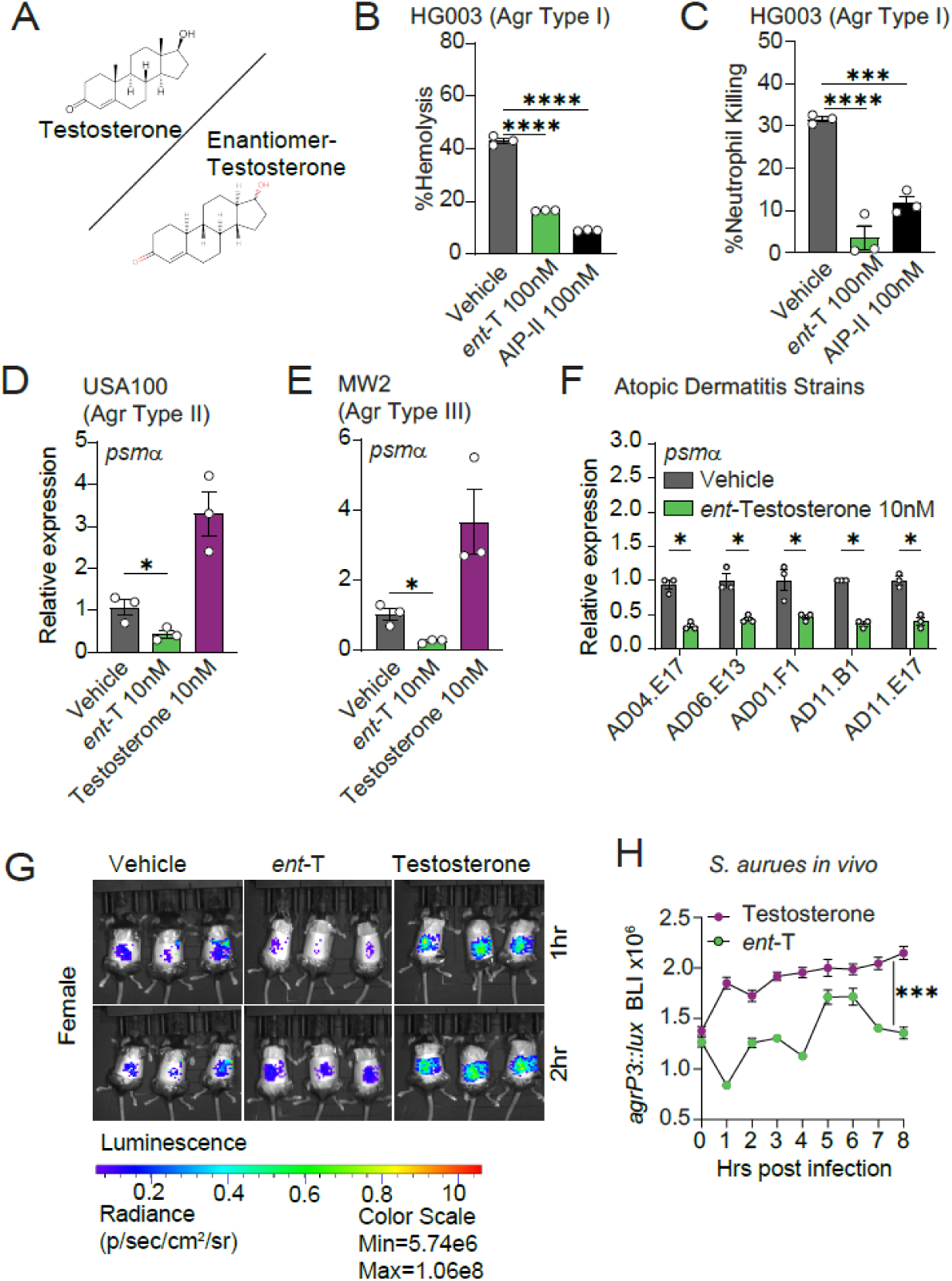
Enantiomer-testosterone inhibits *S. aureus* virulence and *agr* quorum sensing. **(A)**Structure of Testosterone and its stereoisomer enantiomer-Testosterone (*ent*-T). **(B, C)** Wildtype *S. aureus* (HG003) treated with 100nM of *ent*-T, AIP-II, or untreated (vehicle). Percentage of *S. aureus* induced hemolysis (B) and neutrophil killing (C) (*n*=3). **(D-F)** qRT-PCR for *psmα* expression in wild-type *agr* Type II (D), Type III (E), and atopic dermatitis (F) *S. aureus* strains treated with 10nM of testosterone or 10nM of *ent*-T. Relative expression of *psmα* is normalized to housekeeping gene *gyrA* expression. **(G)** Female wild-type mice were epicutaneously infected with 1×10^6^ CFUs *agr-*P3 *S. aureus* reporter treated with testosterone or *ent*-T. Bioluminescence quantified over time. Testosterone (*n*=4), *ent*-T(*n*=4), vehicle (*n*=4). **(H)** Quantification of (G). Means ± SEM (error bars) are plotted.\**p* < 0.05; \*\**p* < 0.01; \*\*\**p* < 0.001, \*\*\*\**p* <0.0001, ns, not significant by two-tailed unpaired *t*-test.

## Discussion

Here we have identified that *S. aureus* pathogenesis is regulated at the skin surface by the secreted androgens, testosterone and DHT. Testosterone regulates *S. aureus* pathogenesis specifically through stimulation of the *agr* quorum sensing system and can signal this system independently of the well characterized bacterially produced AIP. Further, in the absence of host testosterone, *S. aureus* infection is markedly diminished at the skin surface and sex-dependent differences in infection are markedly reduced. Testosterone signaling of *agr* requires the presence of the AgrC histidine kinase and its response regulator AgrA and predictive modeling suggests that testosterone might bind directly to the transmembrane protein, AgrC. Other hormones such as progesterone, estradiol, and pregnenolone, had no impact on quorum sensing in *S. aureus.* A unique feature of both testosterone and DHT are the carbonyl and hydroxyl groups that extended beyond the steroid rings. The hormones that are unable to stimulate *agr* lack both moieties, potentially explaining the ability of testosterone and DHT to activate *agr* signaling and the inability of the other classes of hormones to mediate this pathway.

In contrast to progesterone and estradiol that had a neutral effect on *S. aureus, ent*-T, the stereoisomer of testosterone, decreased quorum sensing and decreased bacterially induced hemolysis and neutrophil killing. *ent*-T also inhibited quorum sensing during *S. aureus* infections *in vivo*. Our findings demonstrate that *S. aureus* responds to testosterone in an enantioselective manner, with testosterone generating activating signals and its chiral counterpart, *ent*-T leading to inhibition of the same phenotypes. Since natural hormones and their enantiomers have been shown to have the same physiochemical properties within membranes(*47*), the distinct phenotypes of testosterone and *ent*-T suggest that the effects of these hormones on *agr* signaling are independent of any predicted impacts of androgen hormones on bacterial membranes.

The skin is known to generate antimicrobial proteins, antimicrobial lipids and nitric oxide that inhibit *S. aureus* pathogenesis (*1, 8, 9, 50–52*). Our work uniquely describes how the skin potentiates the infectious phenotypes of a pathogen at the skin surface through the secretion of testosterone. Indeed, though the biosynthetic mutant had reduced hemolysis and neutrophil killing in *in vitro* assays (Fig. 3B), it retained the ability to sustain *in vivo* skin infections in mice with normal hormone production (Fig. 3G, H). Thus, the host derived hormone signal is sufficient to activate *S. aureus in vivo.* Our work adds to other examples of host hormones and cholesterol-derived molecules that supply activating interkingdom signals to microorganisms with which they have co-evolved (*53–57*). These data provide a novel dissection of host molecules curating the virulence phenotypes of an opportunistic pathogen at the skin surface.

Given their capacity to cue the microbiota, our findings define the need for greater characterization of skin-secreted small molecules and to understand their regulation. Recently, we quantified hormone secretion at the skin surface and illuminated large topographic differences in androgen secretion in young adult humans without skin disease (*17*). We have also identified that IL-4 receptor signaling can regulate androgen production (*18*). More work is needed to determine how androgens may shift in skin conditions mediated by allergic inflammation, such as atopic dermatitis, and to quantify hormonal shifts at the skin surface throughout the human life cycle.

Herein we also show that testosterone regulates *S. aureus* in a dose-dependent manner and that it can limit the ability of other quorum sensing inhibitors to negatively regulate *S. aureus.* Thus, *S. aureus* phenotypes are an aggregate of environmental signals generated from the host and signals generated by the microbiota (*58*). Probiotic approaches that aim to control *S. aureus* virulence through inhibition of the quorum sensing system must take into account the concentration dependent stimulation of *agr* by host derived testosterone and DHT (*39, 59*). Our findings suggest that these therapeutics may be augmented by regulation of host androgen production or exposure to *ent*-T.

## Acknowledgments

Alex Croft, Ph.D. for discussions and aid in experimental design.

## Author contributions

Conceptualization: TAH, MSJ, JK ARH,

Methodology: TAH, MSJ, JK, TS

Investigation: MSJ, MC, TS, MA, MB, RAK, JK

Visualization: MSJ, TAH, TS, RAK, JK,

Funding acquisition: TAH, ARH

Project administration: TAH

Supervision: TAH, ARH

Writing – original draft: TAH, MSJ

Writing – review & editing: TAH, MSJ, MC, MB, TS, JK, ARH

## Data availability

Sequencing data have been submitted to National Center for Biotechnology Information Sequence Read Archive under BioProject accession number PRJNA1071176.

## Material availability

Reagents created for this study including the *Hsd3b6^Δskin^* mice and bioluminescent strains of *S. aureus* are available upon request to Dr. Harris-Tryon.

## Supplementary Materials

### Materials and Methods

#### Mice

Conventionally raised C57BL6/J male and female mice aged 6-9 weeks old mice were purchased from the Jackson Laboratory. C57BL6/J wild-type, *K14^Cre+/-^*(*60*), and *Hsd3b6^fl/fl^* (fig. S2A) mice were bred and maintained in the specific pathogen-free (SPF) barrier facility at the University of Texas Southwestern Medical Center at Dallas. The generation of *Hsd3b6^ΔSkin^* (*K14^Cre+/-^; Hsd3b6^fl/fl^)* is described below. Mice were co-housed with 3–5 mice per cage in all experiments. All mice were housed under a 12-hour-light:12-hour-dark cycle. Mice were fed ad libitum with free access to drinking water according to protocols approved by the Institutional Animal Care and Use Committees (IACUC) of UT Southwestern Medical Center

*Hsd3b6^fl/fl^* (C57BL6/J), with *loxP* sites surrounding the first coding exon of *Hsd3b6* were generated using CRISPR/Cas9 genome editing with guide RNAs targeting regions of the *Hsd3b6* locus (fig. S2A). Guide RNAs were injected into fertilized C57BL/6J embryos by the Children’s research institute mouse genome engineering facility at UT Southwestern. Healthy blastocytes were implanted in pseudo-pregnant mice. The resulting litter was screened by genomic sequencing to detect insertion of *loxP* sites and mice were bred to homozygosity and backcrossed with wild-type C57BL/6 mice. To generate *K14Cre^+/-^;Hsd3b6^fl/fl^(Hsd3b6^ΔSkin^*) mice, *Hsd3b6^fl/fl^* mice were crossed with *K14Cre*^+/-^ mice to generate *K14Cre^+/-^;Hsd3b6^fl/+^*mice, *K14Cre; Hsd3b6^fl/+^* mice were crossed to *Hsd3b6^fl/fl^*mice to obtain experimental mice *K14Cre; Hsd3b6^fl/fl^* (*Hsd3b6^ΔSkin^*) and corresponding controls, *Hsd3b6^fl/fl^*. *Hsd3b6^fl/fl^* and *Hsd3b6^ΔSkin^* status was determined using PCR primers (Table S1) and resolving on a 3% Agarose gel.

#### Quantification of Serum and Skin Hormones in Mouse Samples

Age and sex matched *Hsd3b6^fl/fl^* and *Hsd3b6^ΔSkin^*mice were analyzed. Blood samples were obtained from the retro-orbital vein of anesthetized mice followed by serum isolation with the micro sample tube Serum Gel (SARSTEDT, Cat # 41.1378.005). Skin hormones were quantified from skin secretions. After anesthesia with isoflurane, hair was removed with depilatory cream and shaving. After 24h, Sebutape ® (Clinical and Derm LLC, Texas) was applied to dorsal surface for 15 minutes (fig. S3A). Steroid extraction was performed as previously described (*17, 18, 61, 62*). Sebutape® was removed and placed in 3 ml of chromatography-mass spectrometry grade methanol (Thermo Fisher Scientific, Pittsburg, PA, Cat#A456-500) in an 8 ml polytetrafluorethylene/rubber-lined vial (Thermo Fisher Scientific, Cat#03-343-3E). The sample was then dried by vacuum centrifuge at 40°C and stored at −20°C until analysis. For analysis, samples were reconstituted with 100 μL of kit assay buffer. Steroid Hormone Quantification of progesterone, testosterone, and DHT was measured by mouse specific immunoassay (My BioSource, USA, Cat# MBS7606191, MBS266250, MBS760829) following the manufacturer’s instructions.

#### FACS analysis skin

Ear skin from the epidermis of 8-12 weeks old wildtype, *HSD3B6^fl/fl^* and *HSD3B6^Δskin^*mice were digested with DNase-I at 1-mg/mL (Sigma-Aldrich; DN25-1G), Liberase-TL at 0.32-mg/mL (Sigma-Aldrich; 5401020001), Collagenase-D at 6-mg/mL (Sigma-Aldrich; 11088858001) in RPMI media, minced and incubated for 1 hr at 37°C at 1400 rpm on a thermocycler, and passed through a 70-µm cell separation filter. After washing, the cell pellet was suspended in FACS buffer (PBS with 3% BSA and 2 mM EDTA). The cell suspensions were transferred to a 96 well plate with v-bottom (Corning Inc: Costar; 3894) and processed further. Cell viability was determined with Ghost-Dye-Red-710 (Cytek; SKU 13-0871-T100). To prevent non-specific antibody binding the cells were stained with Fc receptor blocking with anti-mouse CD16/32 antibody (BD Biosciences; 553142). Different cell populations were assessed with the following antibodies re-suspended in FACS buffer for 15 minutes in 4°C. Brilliant-Violet (BV)-650-anti-CD45 (Biolegend; 103151), FITC-anti-CD3 (Cytek; SKU 35-0032-U025), BV-421-anti-CD11b (Biolegend;), BV711-anti-F4/80 (Biolegend;). The gating strategy used to determine cell populations is as follows: For total leucocytes singlet live cells were gated on CD45+ markers, from which the total T cells were identified as CD45+CD3+ cells, macrophages were identified as CD45+CD11b+F4/80+ cells. Cells were acquired by using NovoCyte flow cytometer and analyzed using NovoExpress software.

#### Immunofluorescence microscopy

Mouse skin samples were fixed in formalin and embedded in paraffin by the UT Southwestern histology core. Samples were deparaffined with xylene followed by rehydration with decreasing concentrations of ethanol. Heat induced antigen retrieval was attained in 10 mM sodium citrate buffer. Sections were washed briefly and blocked for an hour in blocking/permeabilization buffer (PBS+5% Goat serum+0.5%Triton X-100). Sections were incubated in blocking/permeabilization buffer overnight with the following antibodies: anti-HSD3B6 (2.5 µg/mL; Biorbyt; orb592071), anti-cytokeratin-14 (1 µg/mL; Santa Cruz Biotechnology; sc-53253). After a brief wash in PBST (PBS+0.2% Tween-20) sections were incubated with corresponding secondary antibodies: Donkey anti-rabbit alexa-fluor-647 (2 µg/mL; Jackson Immuno; 711-605-152), Donkey anti-mouse alexa-fluor-594 (2 µg/mL; Thermofisher Scientific; A-21203). The slides were then washed briefly in PBST and mounted with 4, 6-diamidino-2-phenylindole (DAPI) containing mounting medium (SouthernBiotech; 0100-20). Images were processed using ZEISS 780 confocal microscope.

#### TEWL measurement

Transepidermal water loss (TEWL), a measure of barrier function and integrity, of mice dorsal skin was measured using Vapometer (Delfin Technologies) according to manufacturer instructions(*63*).

#### Bacterial strains and plasmids

*S*. *aureus* strains (Table S1) were streaked on tryptic soy agar (TSA) plates and grown overnight at 37 °C. Single colonies were selected and cultured in TSB at 150 RPM at 37 °C in a shaking incubator overnight followed by a 1:100 subculture at 37 °C in a shaking incubator to obtain mid-logarithmic phase bacteria. For fusion reporter strains, all *in vitro* cultures were performed using TSB in the presence of 10 µg/mL of chloramphenicol. Bacteria were pelleted, washed, and re-suspended in either TSB for *in vitro* experiments or PBS for *in vivo* experiments. HG003, *ΔagrC, ΔagrBD,* and *ΔagrA* mutant strains were obtained from Dr. Ferric Fang (*9*). *S. aureus* strains from atopic dermatitis skin obtained from Drs. Julie Segre and Heidi Kong (*25*). All other strains from the collections of the Horswill and the Harris-Tryon labs.

#### Construction of a *lux* expressing *S. aureus* strain

As described previously(*64*), the integrated *luxCDABEG* cassette was transduced into *S. aureus* strains HG003, *ΔagrBD* and *ΔagrC* obtained from the lab of Ferric Fang (*9*) using phage 11 generating strains AH6222 (*lux+*), AH6224 (*lux+*) and AH6223 (*lux+*), respectively.

#### *In vitro* luminescence assays

*S. aureus* strains, HG003, *ΔagrBD* and *ΔagrC* expressing Lux (φ11::LL29*luxCDABEG*) and quorum sensing lux (pAmiAgrP3lux) plasmids (HG003 (AH6225) *agr* Type I, USA100 (AH430). Type II and MW2 (AH1747) *agr* Type III) (*52, 65, 66*) were grown in TSB supplemented with antibiotic selection and subcultured in TSB 1:200 into fresh TSB containing steroid hormone. Assay completed in Opaque-sided, 96-well, clear bottom, tissue-culture treated plates with a final well volume of 200 μL. Bioluminescent signals (photons/0.1 second acquisition time) were measured by BioTek H1 Synergy plate reader. Experiments completed in triplicate, with *agr* type specific AIPs AIP-I (Peptide Institute, Inc., Cat# 4515-v), AIP-II (Peptide Institute, Inc., Cat# 4516-v), and AIP-III(Peptide Institute, Inc., Cat# 4517-v) as positive control.

#### Hemolysis assay

Hemolysis assay completed as previously described (*67*) with the following modifications. Overnight cultures of HG003, *ΔagrBD*, and *ΔagrC* strains were inoculated 1:200 into 10 mL of TSB containing testosterone, AIP-I, or vehicle alone at concentrations of 10 nM. Cells were grown to mid-log phase (OD600 nm 0.6). Supernatant of 1ml of culture was filter sterilized using Millex® sterile syringe filters of 0.22 µm pore size (Cat#SLGV033RS). Filtered supernatant diluted 1:1 with PBS was added to 25 µL of human blood to a 96-well V-bottom plate and incubated with agitation at 37°C for 1 hr. After spinning 1000RPM for 10 min, the supernatant was transferred to a flat-bottom 96-well plate. Absorbance read at 541 nm for hemoglobin using a BioTek H1 Synergy plate reader. % hemolysis calculated using the following formula: (A541 of RBC treated sample-A541 of buffer)/ (A541 of H20-A541 of buffer) Buffer (PBS) = baseline, H20 = 100% hemolysis.

#### Neutrophil Killing assay

*S. aureus* induced neutrophil killing measured as previously described (*68*). HG003, *ΔagrBD* and *ΔagrC* strains were treated with testosterone, AIP-I or vehicle at concentrations of 10nM and allowed to grow to mid log phase (OD600nm 0.6). Purified Human Neutrophils (IQ Biosciences) were seeded at 1 × 10^5^ cells per well into 96-well plate in 90 μL of RPMI. 10 μL of bacterial supernatants were added (final concentration of 10%). After 3h incubation at 37°C, 5% CO2, the plates were centrifuged at 250g, 10 min, and resulting supernatants were used to measure lactate dehydrogenase (LDH) leakage from damaged cells as the marker of neutrophil lysis with an LDH Cytotoxicity Detection Kit (Invitrogen, Cat# 2570393). Percent neutrophil lysis was calculated using neutrophils incubated with 10% of RPMI as zero percent lysis control, and neutrophils incubated with 0.2% Triton X-100 defined as 100 percent lysis.

#### Quantitative real-time PCR

HG003, USA100 (AH3684), MW2 (AH843), *ΔagrBD, ΔagrA*, *ΔagrC* and atopic dermatitis strains were treated with testosterone and/or respective AIPs at concentrations of 10nM and allowed to grow to mid-log phase (OD600nm 0.6). Cells were pelleted and lysed with lysis matrix B tubes containing 0.1mm silica spheres (MP Lysing Matrix Tubes, Cat#174701) and lysostaphin (Sigma, Cat# L7386) at room temperature, and RNA was purified using the RNeasy Mini Kit (Qiagen, Cat# 74104). RNA was quantified by absorbance at 260 nm, and its purity was evaluated by the ratios of absorbance at 260/280 nm. RNA was used as a template to generate cDNA with the High-Capacity Reverse Transcription Kit (Applied Biosystems, Cat#01071619). Quantitative real-time PCR was performed by amplifying cDNA with Power SYBR Green Master Mix (Applied Biosystems, Cat# 2749999) and QuantStudio 7 Flex Real-Time PCR System (Applied Biosystems). Relative expression values were calculated using the comparative Ct (ΔΔCt) method, and transcript abundances were normalized to *gyrA* transcript abundance. The primer sequences are shown in Table S2.

#### RNA Seq

RNAseq was performed as previously described (*69*). Briefly, cultures of HG003 were grown in TSB with 10nM testosterone, pregnenolone, or DMSO alone in triplicate to an optical density of 0.6 at OD600 nm. Cells were harvested and treated with RNA Protect Bacteria Reagent (Qiagen, Cat# 76526). Cells were lysed using lysostaphin (Sigma, Cat# L7386) and RNA purified using the RNeasy mini kit (Qiagen, Cat# 74104) and sample quality was affirmed via Bio analyzer (Agilent). Ribosomal RNA was depleted using RiboCop for bacterial META Removal Kit (Lexogen). cDNA libraries were generated at the University of Michigan Microbiome core using the CORALL RNA-seq Library Prep Kit (Lexogen). Samples were barcoded, pooled and sequenced in 125×125 paired-end reads on an Illumina HiSeq 2000 sequencer. Raw sequencing reads in fastq format were aligned and annotated to the *S. aureus* NCTC8325 reference genome with annotated sRNA(*70*) using QiagenCLC Genomics Workbench default settings (version 21.0.5): mismatch cost, 2; insertion and deletion cost, 3; length and similarity fraction, 0.8. Normalization and differential expression calculations of uniquely mapped bacterial transcripts were performed using CLC. All transcripts with an FDR adjusted *p*-value <0.05 were considered significant.

#### *S. aureus* skin infections

Prior to mouse infection studies, mice were acclimatized to the animal biosafety level 2 (ABSL-2) animal housing facility. Age, strain and sex matched C57BL/6 male and female mice, *Hsd3b6^fl/fl^*and *Hsd3b6^ΔSkin^* were used in the study. A previously described mouse model of epicutaneous *S. aureus* exposure was followed (*21, 50*). Briefly, the dorsal skin of anesthetized mice (2% isoflurane) were shaved and depilated (Nair cream). After 24 hours, bioluminescent *S. aureus* strains were grown to mid-log-phase, pelleted and resuspended in PBS to achieve inoculum containing 1×10^6^ CFU. A 100 μL volume of PBS containing 1×10^6^ CFU with or without 10nmoles of testosterone, AIP-I, or the same volume of vehicle was placed on a sterile gauze pad and attached to the shaved skin with transparent bio-occlusive dressing (Tegaderm; 3M, Henry Schein medicals, Cat#1622W), and secured with adhesive bandages (BAND-AID, Johnson and Johnson, American white cross, Cat#1275033) for 4 days. Photons emitted from luminescent bacteria were collected during an auto exposure using the IVIS Lumina3 imager machine and living image software (Xenogen, Alameda, CA). Bioluminescent image data are presented on a pseudocolor scale (blue representing least intense and red representing the most intense signal) overlaid onto a gray-scale photographic image. Using the image analysis tools in living image software, circular analysis windows (of uniform area) were overlaid onto dorsal regions of infection area, and the corresponding bioluminescence values (total flux) were measured and plotted versus days after infection. Mice were randomly assigned to treatment groups, and at experimental endpoints, mice were humanely euthanized using carbon dioxide inhalation.

#### Measurement of quorum sensing *in vivo*

*S. aureus* strains expressing quorum sensing lux (pAmiAgrP3lux) plasmids as described above (*30*) were grown in TSB medium containing chloramphenicol overnight at 37°C in a shaking incubator set to 150 rpm. Overnight cultures were diluted 1:100 TSB with chloramphenicol to mid-logarithmic phase and then pelleted and washed twice in PBS and resuspended in sterile saline. 100 μL of PBS inoculum suspensions containing 1×10^6^ CFUs were placed on a sterile gauze pad (1×1cm) and attached to the shaved skin with transparent bio-occlusive dressing, with or without testosterone, enantiomer-testosterone, AIP-I, or vehicle (Tegaderm; 3M), and secured with 2 layers of adhesive bandages (BAND-AID, Johnson and Johnson). Beginning immediately after infection, mice were imaged under isoflurane inhalation anesthesia (2%) and continued to take images for every 1hr. Photons emitted from luminescent bacteria were collected during auto exposure using the IVIS Lumina3 imager machine and living image software (Xenogen, Alameda, CA). Corresponding bioluminescence values (total flux) were measured and plotted versus time after infection.

#### In silico docking

The dimeric structure of AgrC was predicted using Alphafold2 (*42, 43*) as implemented in Colab (*42, 43*) and then AIP-I was *in silico* docked using SWISSDOCK (*44–46*) onto the sensory domain of a single subunit, consisting residues 1 through 207, using NMR derived coordinates for AIP-I (*44–46*). The AIP-I docking solution that appeared most consistent with the structure-activity relationships reviewed in Thoendel *et al*. (*5, 6*) was selected as the target AgrC-AIP-I complex for docking of steroids, using stereospecific compound templates from PubChem; testosterone (CID 6013). All visualization of *in silico* results was done using PyMOL [ver 2.5.2, Schrödinger, LLC].

#### Quantification and Statistical Analysis

Statistical details of experiments can be found in the figure legends, including how significance was defined and the statistical methods used. Data represent mean ± standard error of the mean. Formal randomization techniques were not used; however, mice were allocated to experiments randomly and samples were processed in an arbitrary order. Mouse skin samples that were determined to be in the anagen hair cycle were excluded. All statistical analyses were performed with GraphPad Prism software, except the bioluminescent imaging data that was analyzed as described above. To assess the statistical significance of the difference between two treatments, we used two-tailed Student’s *t*-tests. To assess the statistical significance of differences between more than two treatments, we used one-way ANOVA. Outliers within experiments were identified by Grubb’s test and removed. For the RNAseq experiments, expression data was analyzed with CLC. All transcripts with an FDR adjusted *p*-value <0.05 were considered significant.

---

**Fig. S1.**
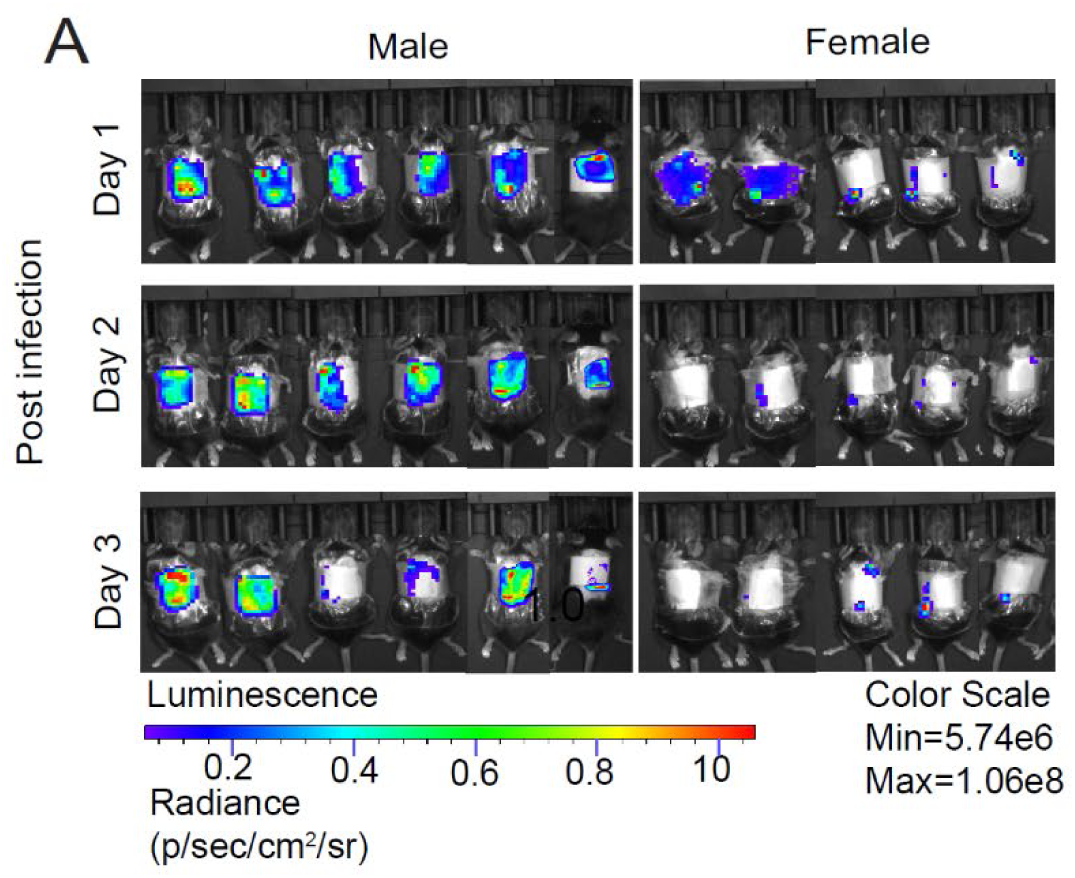
Sex bias of *S. aureus* skin infection. **(A, B)** Wildtype male mice (*n*=6) and female mice (*n*=7) were epicutaneously challenged with 1×10^6^ CFUs with bioluminescent MRSA (MRSA:*:lux*) for 3 days and bioluminescence quantified over time. **(A)** *In vivo* Bioluminescence images.

**Fig. S2.**
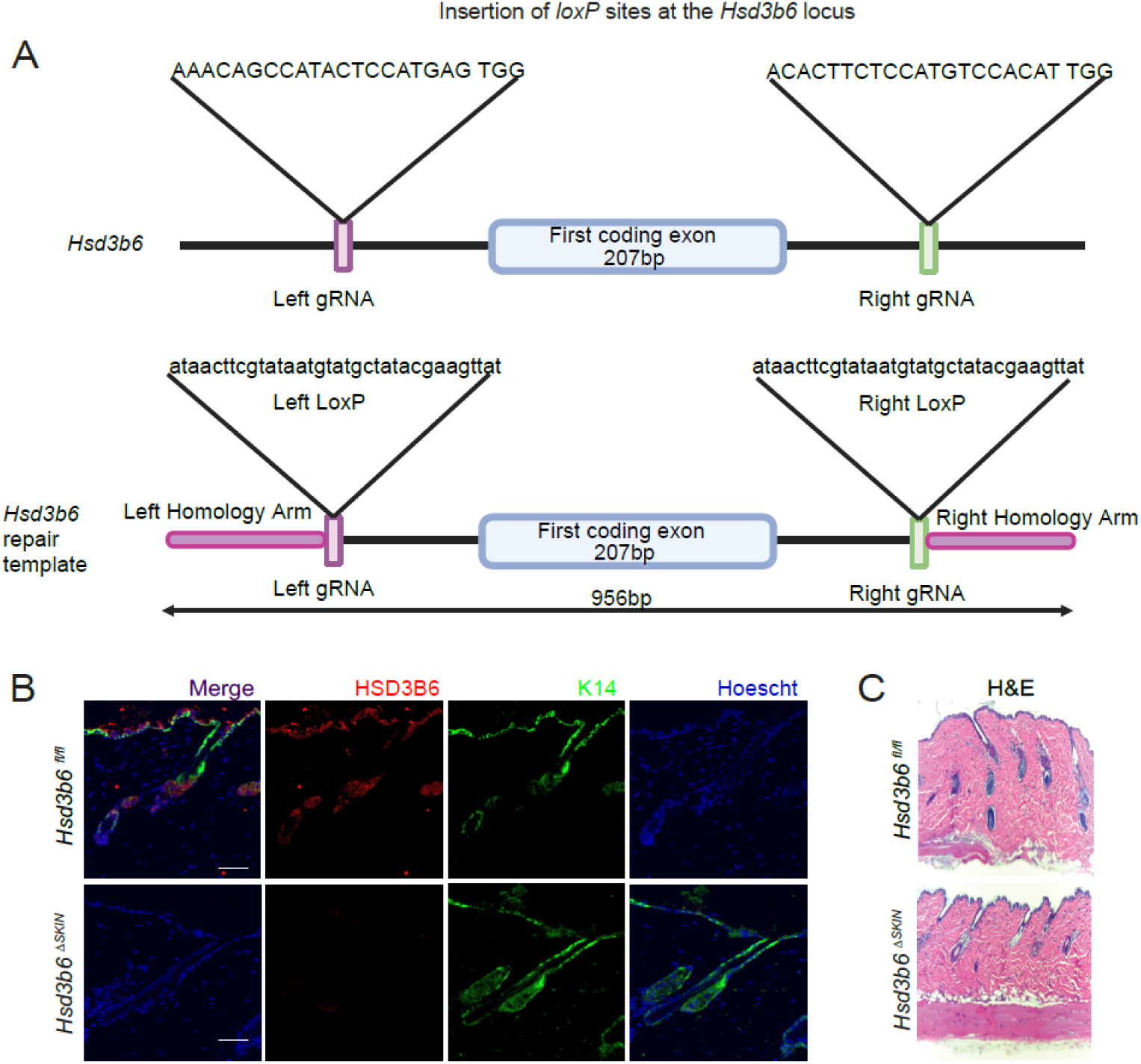
Generation and validation of *Hsd3b6^Δskin^* mice by CRISPR/Cas9 genomic targeting. **(A)** Schematic diagram of CRISPR/Cas9-mediated gene insertion of *loxP* sites flanking the first coding exon of the *Hsd3b6* locus. **(B)** Immunofluorescence staining of HSD3B6 expression in *Hsd3b6^fl/fl^*and *Hsd3b6^Δskin^* mice skin. Scale 50µM. **(C)** Hematoxylin and eosin staining of *Hsd3b6^fl/fl^*and *Hsd3b6^Δskin^* skin.

**Fig. S3.**
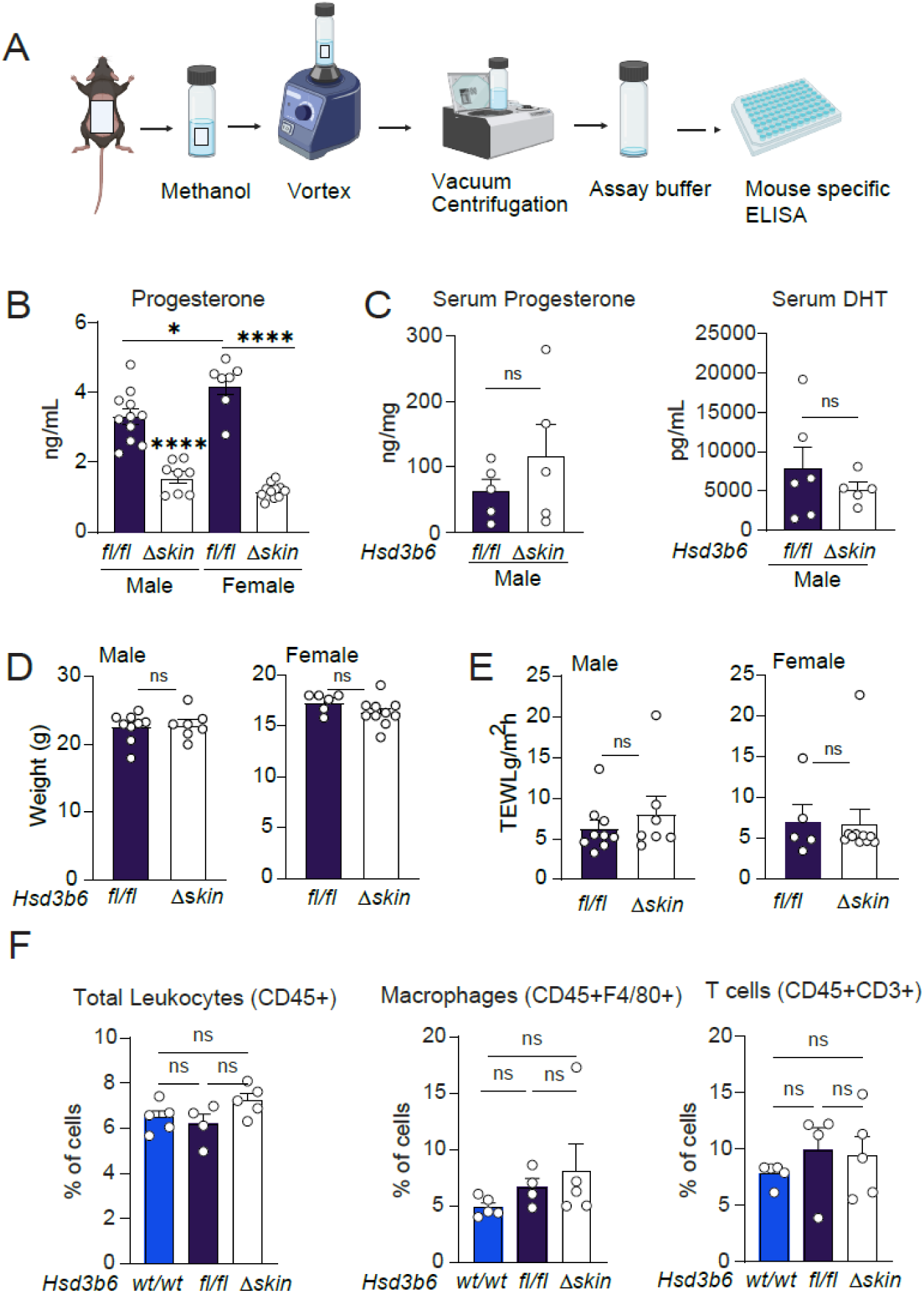
Analysis of the *Hsd3b6^Δskin^* mice compared to controls. **(A)** Skin secretions captured with commercial Sebutape® applied to the dorsal skin after hair removal. Each hormone quantified by hormone specific immunoassay. **(B)** Skin secreted progesterone quantified by immunoassay of male and female *Hsd3b6^fl/fl^ and Hsd3b6^Δskin^* mice. **(C)** Serum progesterone and DHT quantified by hormone immunoassay of male and female *Hsd3b6^fl/fl^*and *Hsd3b6^Δskin^* mice. **(D)** Weight measurements of male and female *Hsd3b6^fl/fl^* and *Hsd3b6^Δskin^* mice. **(E)** Transepidermal water loss (TEWL), a measures of skin barrier integrity, of male and female *Hsd3b6^fl/fl^* and *Hsd3b6^Δskin^* mice measured by Vapometer device. **(F)** Flow cytometry for macrophages, total leucocytes, and T-cells (markers as shown), in the skin of *Hsd3b6^fl/fl^* and *Hsd3b6^Δskin^* mice. *n* as shown. Means ± SEM (error bars) are plotted.\**p* < 0.05; \*\**p* < 0.01; \*\*\**p* < 0.001, \*\*\*\**p* <0.0001, ns, not significant by two-tailed unpaired *t*-test.

**Fig. S4.**
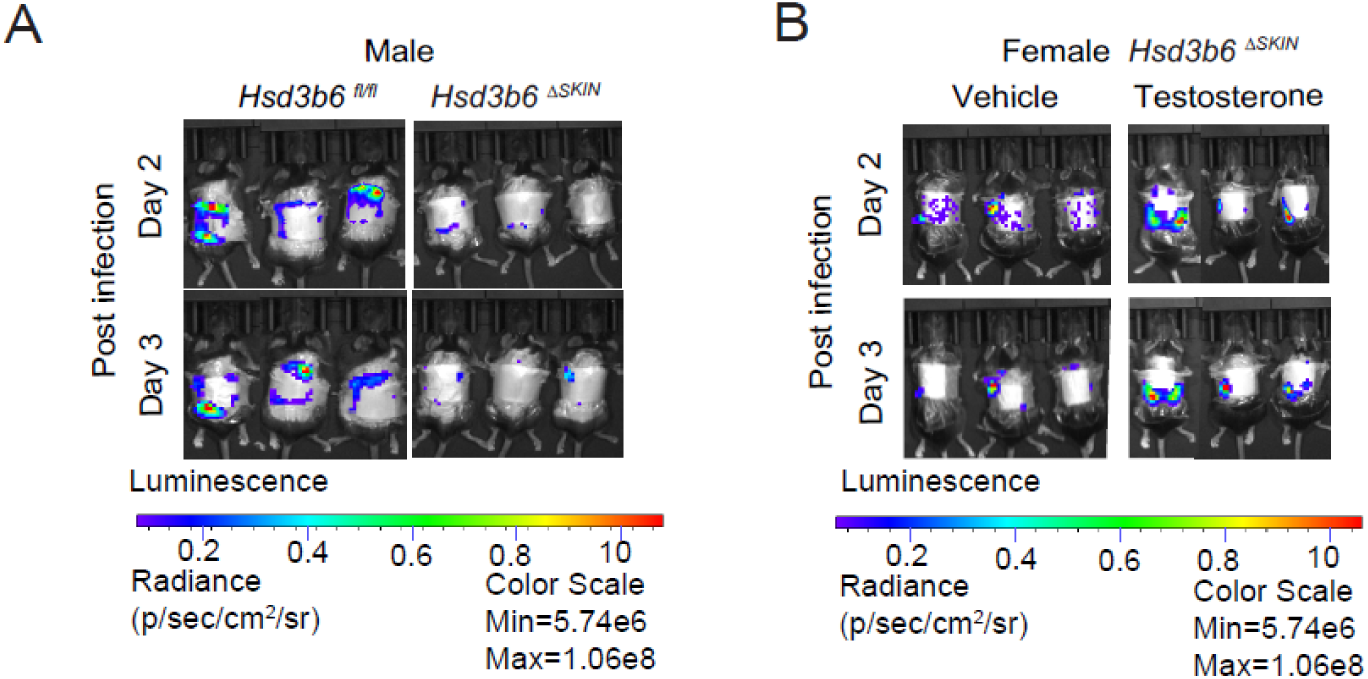
*Hsd3b6^Δskin^* mice are resistant to skin infection with S. *aureus*. **(A)** Male Hsd3b6*^fl/fl^*(*n*=7) and *Hsd3b6^Δskin^* (*n*=7) mice were epicutaneously challenged with 1×10^6^ CFUs with bioluminescent MRSA (MRSA::lux) for 3 days and bioluminescence quantified over time. **(B)** Female *Hsd3b6^Δskin^* mice were epicutaneously challenged with 1×10^6^ CFUs with bioluminescent MRSA (MRSA::*lux*) for 3 days with or without testosterone(*n*=5) and vehicle (*n*=5).

**Fig. S5.**
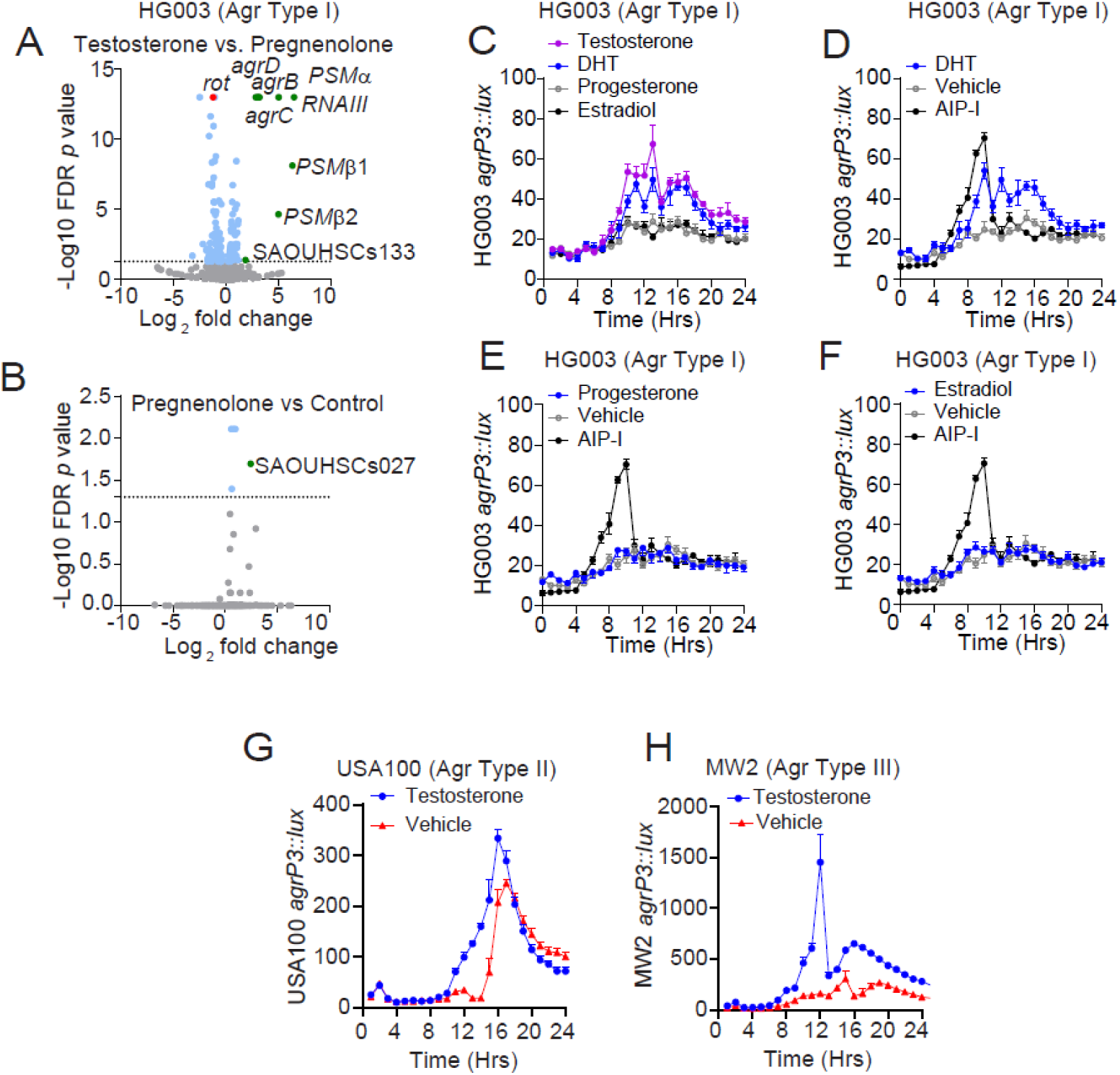
Testosterone and DHT activate quorum sensing readouts in *S. aureus*, but other hormone classes do not activate quorum sensing. **(A, B)** Transcriptomics (RNA-seq) of HG003 strain of *S. aureus* treated with 10nM testosterone, pregnenolone, or untreated with vehicle alone. (A) Volcano plot demonstrating genes with >4-fold change in expression in testosterone transcriptome compared to pregnenolone treated *S. aureus,* (*n*=3). (B) Volcano plot comparing the transcriptome of *S. aureus* treated with pregnenolone compared to vehicle control (*n*=3). **(C-F)** *In vitro* bioluminescence of *agr* reporter (HG003 *agrP3*::*lux*) after treatment with 10nM of testosterone (C), DHT (C, D), progesterone (C, E), AIP-I (C-F), or estradiol(C, F). **(G, H)** *In vitro* bioluminescence of Type II (USA100 *agrP3::lux*) (G) and Type III (MW2 *agrP3::lux*) (H) *agr* reporters treated with 10nM of testosterone (*n*=3).

**Fig. S6.**
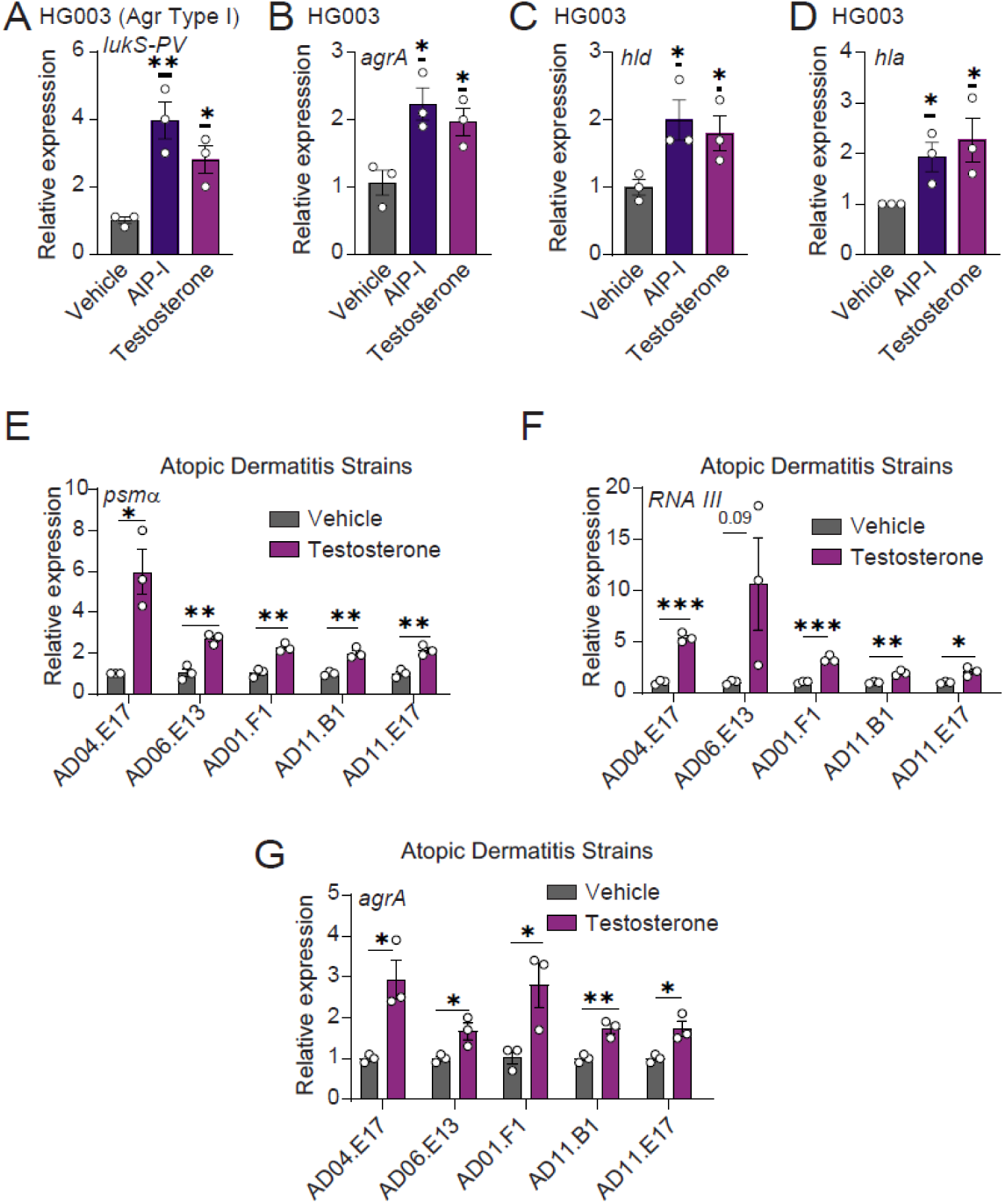
Testosterone stimulates the expression of virulence factors across strains of *S. aureus.* **(A-D)** qRT-PCR of *S. aureus agr* Type I strain HG003 treated with 10nM Testosterone or AIP-I (*n*=3) till mid-exponential growth. Expression of target genes, *lukS-PV*, *hla*, *hld*, and *agrA* normalized to *gyrA* expression. **(E-G)** qRT-PCR of strains of *S. aureus* obtained from patients with atopic dermatitis (*25*) and treated with 10nM of testosterone to the mid-exponential growth (*n*=3). Expression of target gene, *psmα* (E), *RNAIII* (F), and *agrA* (G), normalized to housekeeping gene *gyrA* expression. Means ± SEM (error bars) are plotted.\**p* < 0.05; \*\**p* < 0.01; \*\*\**p* < 0.001, \*\*\*\**p* <0.0001, ns, not significant by two-tailed unpaired *t*-test.

**Fig. S7.**
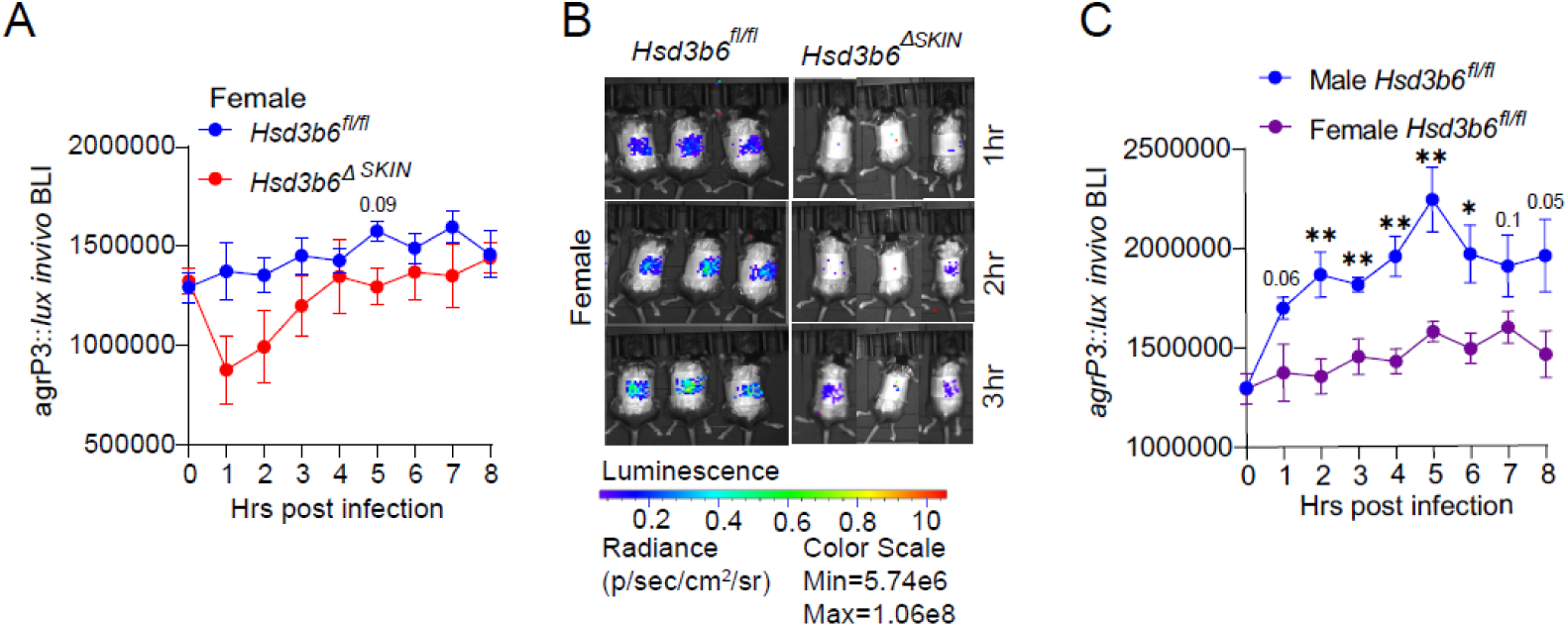
Sexual dimorphism of *agr* quorum sensing in vivo. **(A-C)** In vivo analysis of *agr-*P3 reporter. Female *Hsd3b6^fl/fl^* and *Hsd3b6^Δskin^* mice were epicuataneously infected with 1×10^6^ CFUs of the *agr-P3lux* and bioluminescence quantified over time. (A) Kinetics and (B) representative images. (C) Comparison between male (Fig 2H) and female *Hsd3b6^fl/fl^* mice. Aggregate of two experiments, *n*=5 male and *n*=5 female mice. Means ± SEM (error bars) are plotted.\**p* < 0.05; \*\**p*< 0.01; \*\*\**p* < 0.001, \*\*\*\**p* <0.0001, ns, not significant by two-tailed unpaired *t*-test.

**Fig. S8.**
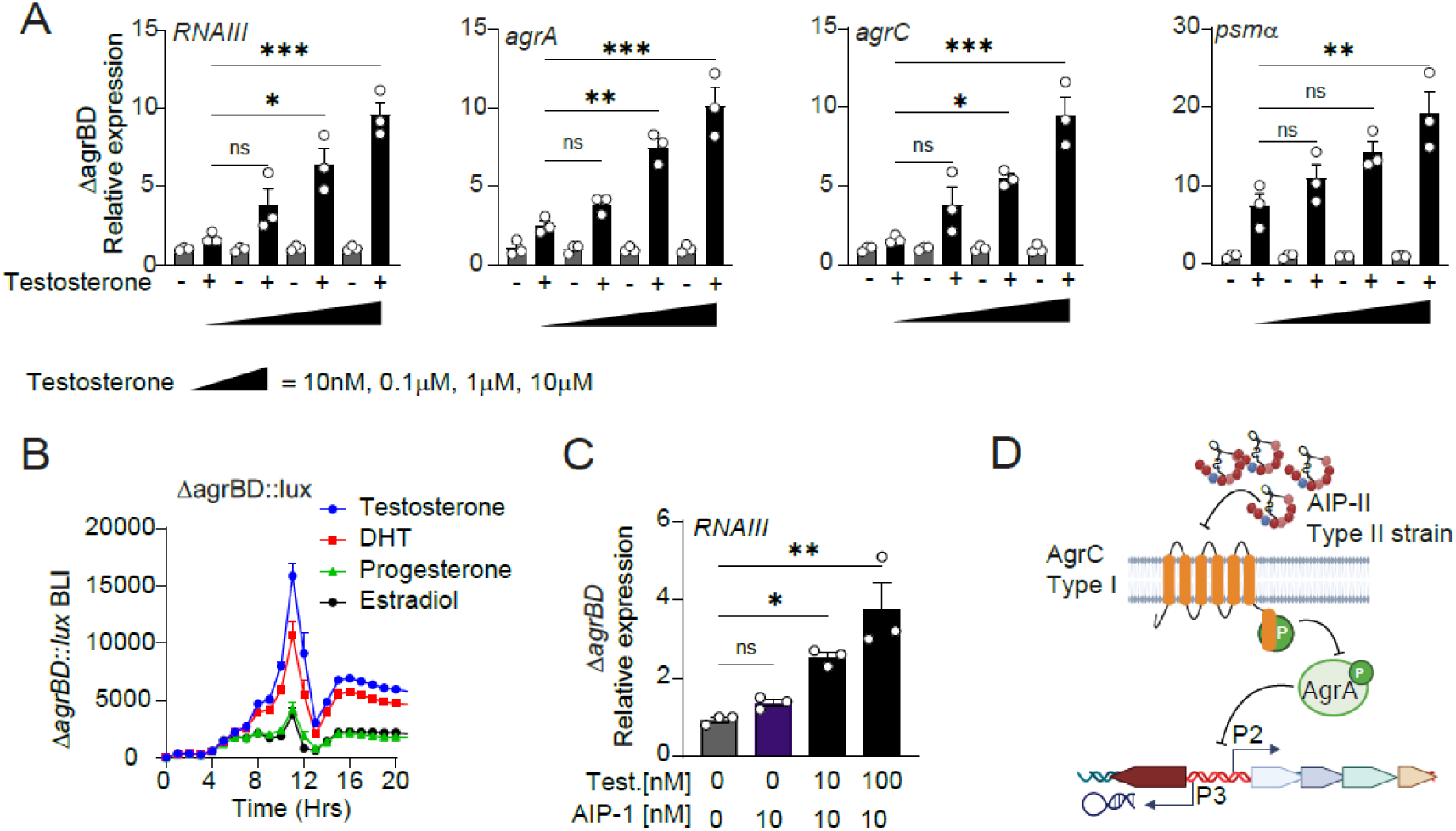
Testosterone stimulates *agr* quorum sensing in *S. aureus* in a dose dependent manner, independent of the autoinducing peptides (AIP). **(A)***S. aureus* biosynthetic mutant (*ΔagrBD*) treated with increasing doses of testosterone, followed by qRT-PCR. *RNAIII, agrA, agrC,* and *psmα* expression normalized to *gyrA* expression. **(B)** *S. aureus* biosynthetic mutant reporter (*ΔagrBD::lux*) treated with 10nM testosterone, DHT, progesterone, or estradiol. **(C)** qRT-PCR for *RNAIII* expression in the biosynthetic mutant strain treated with 10nM AIP-I and 10 nM or 100nM or testosterone. **(D)** Schematic diagram showing inhibitory action of non-cognate AIPs on *agr* signaling. Means ± SEM (error bars) are plotted.\**p* < 0.05; \*\**p* < 0.01; \*\*\**p* < 0.001, \*\*\*\**p* <0.0001, ns, not significant by one-way ANOVA.

**Fig. S9.**
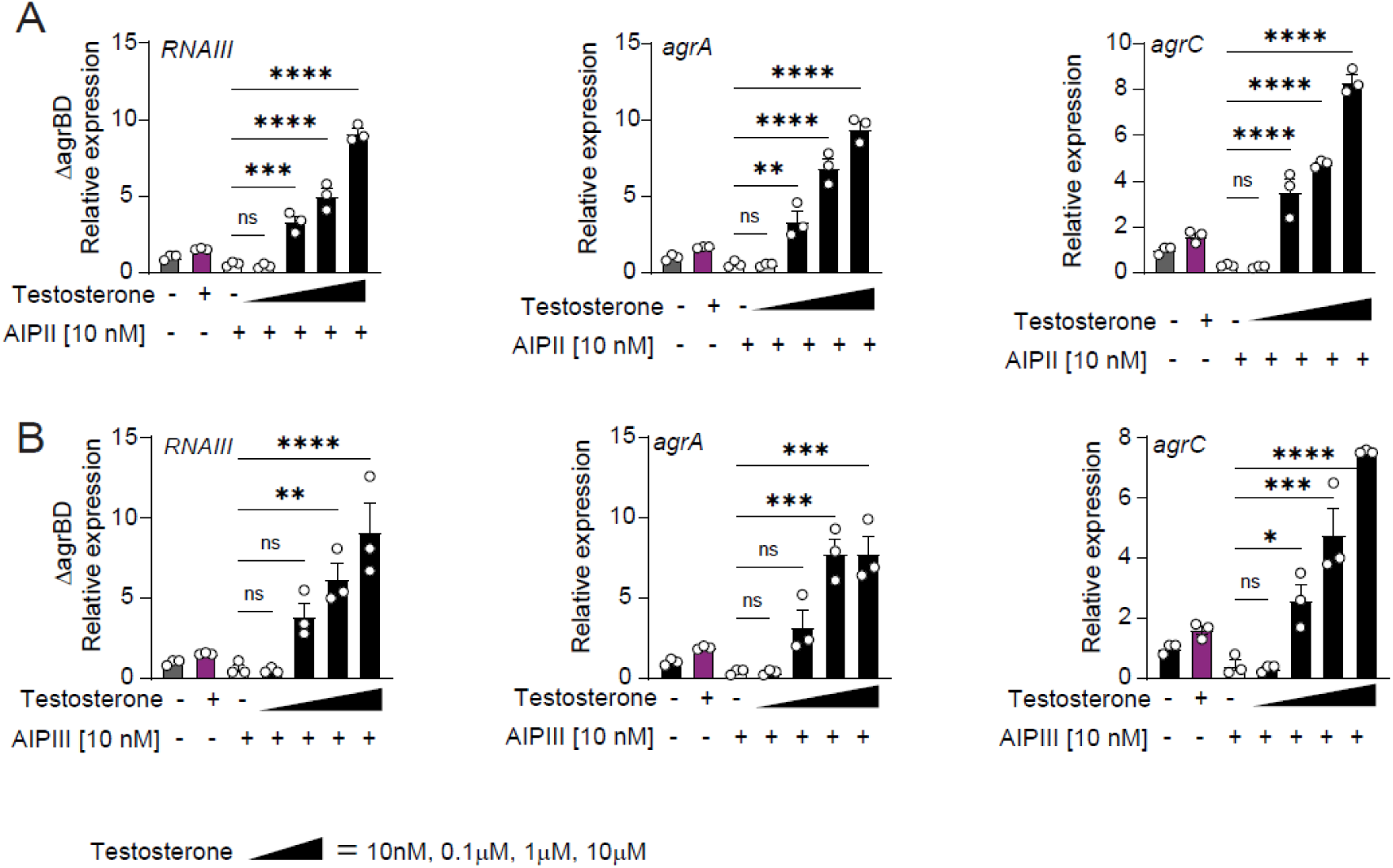
Testosterone can overcome the inhibition of AgrC signaling in a dose dependent manner. **(A, B)** qRT-PCR for *RNAIII, agrA* and *agrC* expression in the biosynthetic mutant, *ΔagrBD*, treated with AIP-II **(A)** or AIP-III **(B)** alone or in combination with increasing concentrations of testosterone (10nM to 10µM). Target gene expression normalized to housekeeping gene *gyrA*. Means ± SEM (error bars) are plotted.\**p* < 0.05; \*\**p* < 0.01; \*\*\**p* < 0.001, \*\*\*\**p* <0.0001, ns, not significant by one-way ANOVA.

**Fig. S10.**
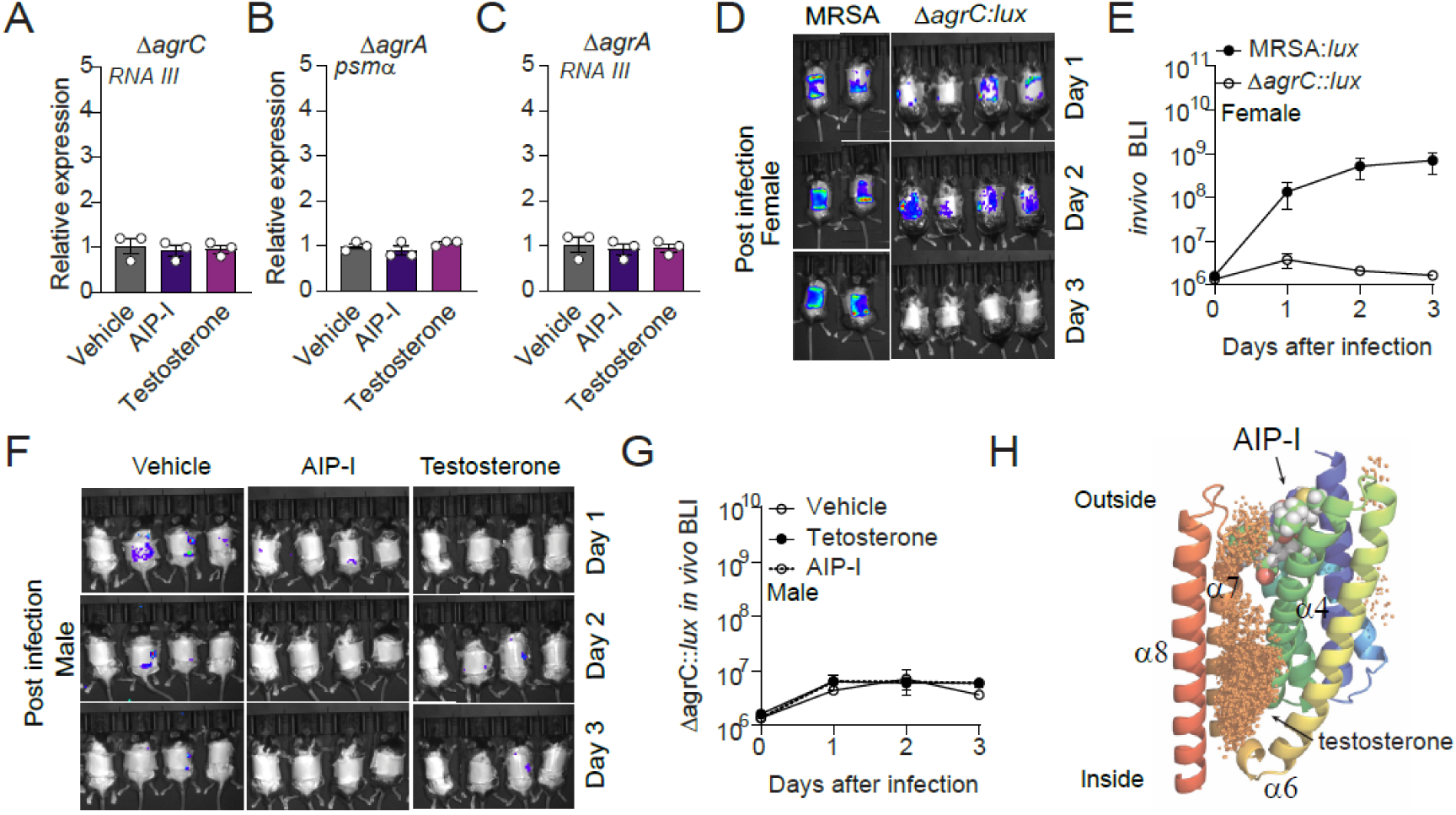
Testosterone stimulation requires *agrC in vivo and in vitro.* **(A-C)** *S. aureus ΔagrC* (A) and *ΔagrA* (B, C) mutant strains were treated with 10nM Testosterone, AIP-I, or untreated (vehicle). *Psmα (B)* and *RNAIII* (C) expression quantified by qRT-PCR and normalized to *gyrA* expression. **(D, E)** Female wild-type mice were epicutaneoulsy infected with 1×10^6^ CFUs of wild-type (MRSA::*lux*) (*n*=5) or agrC histidine kinase deficient (*ΔagrC::lux*) (*n*=6) bioluminescent reporter strain of *S. aureus* and bioluminescence quantified over time. Bioluminescence images (D) and infection kinetics from an aggregate of two experiments (E). **(F, G)** Male wildtype mice were infected for 3 days with 1×10^6^ CFUs of *ΔagrC::lux* treated with testosterone (*n*=5), AIP-I (*n*=5) or vehicle (*n*=5) control. Bioluminescence images (F) and infection kinetics (G). Means ± SEM (error bars) are plotted. ns, not significant by two-tailed unpaired *t-*test. **(H)** Ribbon diagram of the AgrC sensory domain (Alphafold 2) with docking of space filling models of AIP-I and predictive locations of testosterone docking.

**Fig. S11.**
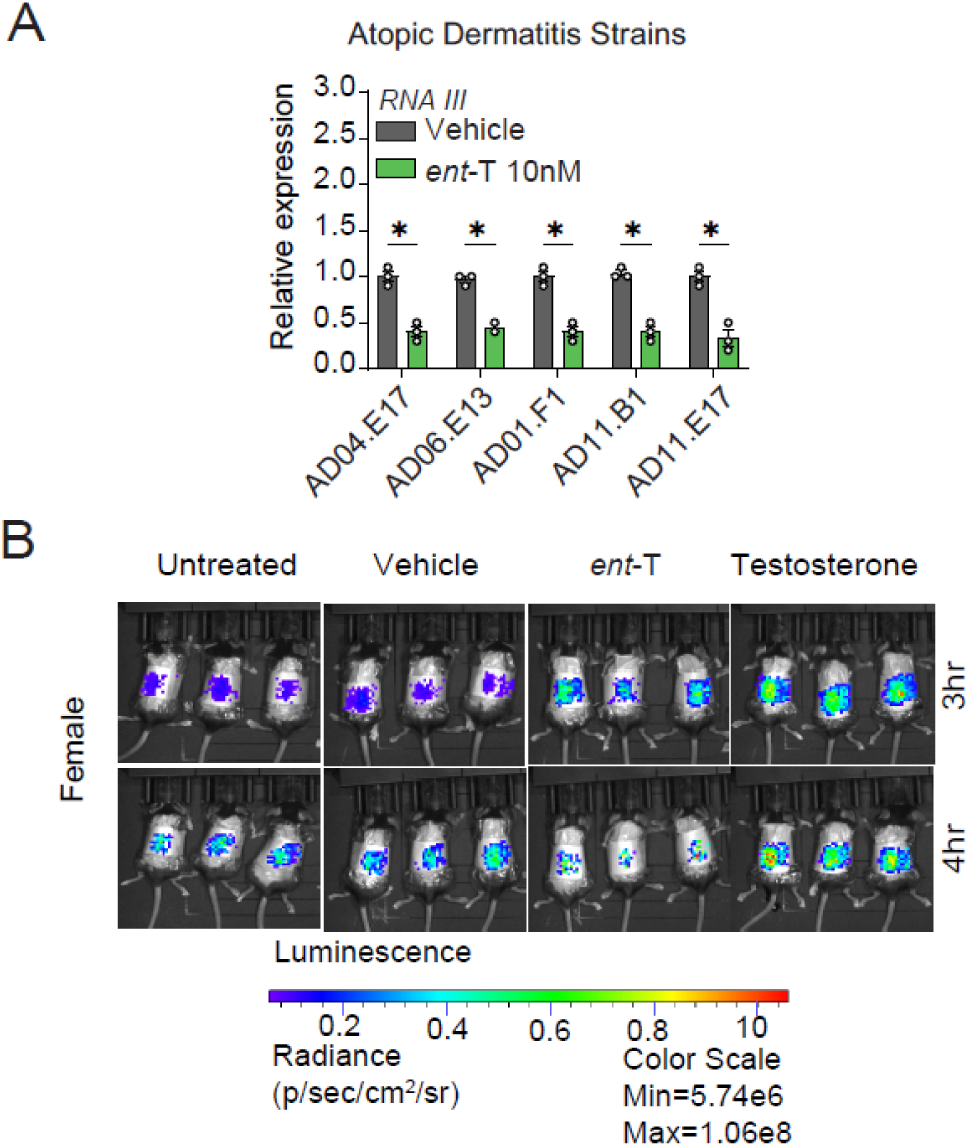
Enantiomer-testosterone (*ent*-T) inhibits *S. aureus* virulence factor expression and quorum sensing. **(A)** qRT-PCR of strains of *S. aureus* obtained from patients with atopic dermatitis (*25*) and treated with 10nM or *ent*-T, the stereoisomer of testosterone (n=3). Expression of target gene *RNAIII* normalized to housekeeping gene *gyrA* expression. Means ± SEM (error bars) are plotted.\**p* < 0.05 by two-tailed unpaired *t*-test. **(B)** Representative bioluminescence images of Fig. 5H, female wild-type mice epicutaneously infected with 1×10^6^ CFUs *agr-*P3 *S. aureus* reporter treated with testosterone, *ent*-T, vehicle, or untreated. (*n*=*4*).

**Supplemental Table 1.** *Staphylococcus aureus* strains used in this work.

| *S. aureus* strains | Reference | Identifier |
| --- | --- | --- |
| HG003 | (9) | RU083 |
| USA100 | (29) | AH3684 |
| MW2 | (29) | AH843 |
| HG003-ΔagrA | (29) | RU111 |
| HG003-ΔagrC | (29) | RU141 |
| HG003-ΔagrBD | (29) | RU299 |
| MRSA SAP430::*lux*ABCDE | (21) | *Staphylococcus aureus*<br>USA300 LAC::*lux* |
| HG003, ϕ11::*LL29luxCDABEG* | This work | AH6222 |
| HG003-ΔagrBD, ϕ11::*LL29luxCDABEG* | This work | AH6224 |
| HG003-ΔagrC, ϕ11::*LL29luxCDABEG* | This work | AH6223 |
| USA300 LAC + pAmiAgrP3*lux* (camR) | (29) | AH2759 |
| HG003+ pAmiAgrP3*lux* (camR) | This work | AH6225 |
| USA100 IA116 + pAmiAgrP3*lux* (camR) | (29) | AH4390 |
| MW2+ pAmiAgrP3*lux* (camR) | (65) | AH3185 |
| HG003-ΔagrBD + pAmiAgrP3*lux* (camR) | This work | AH6227 |
| HG003-ΔagrC+ pAmiAgrP3*lux* (camR) | This work | AH6226 |

**Supplemental Table 2.**
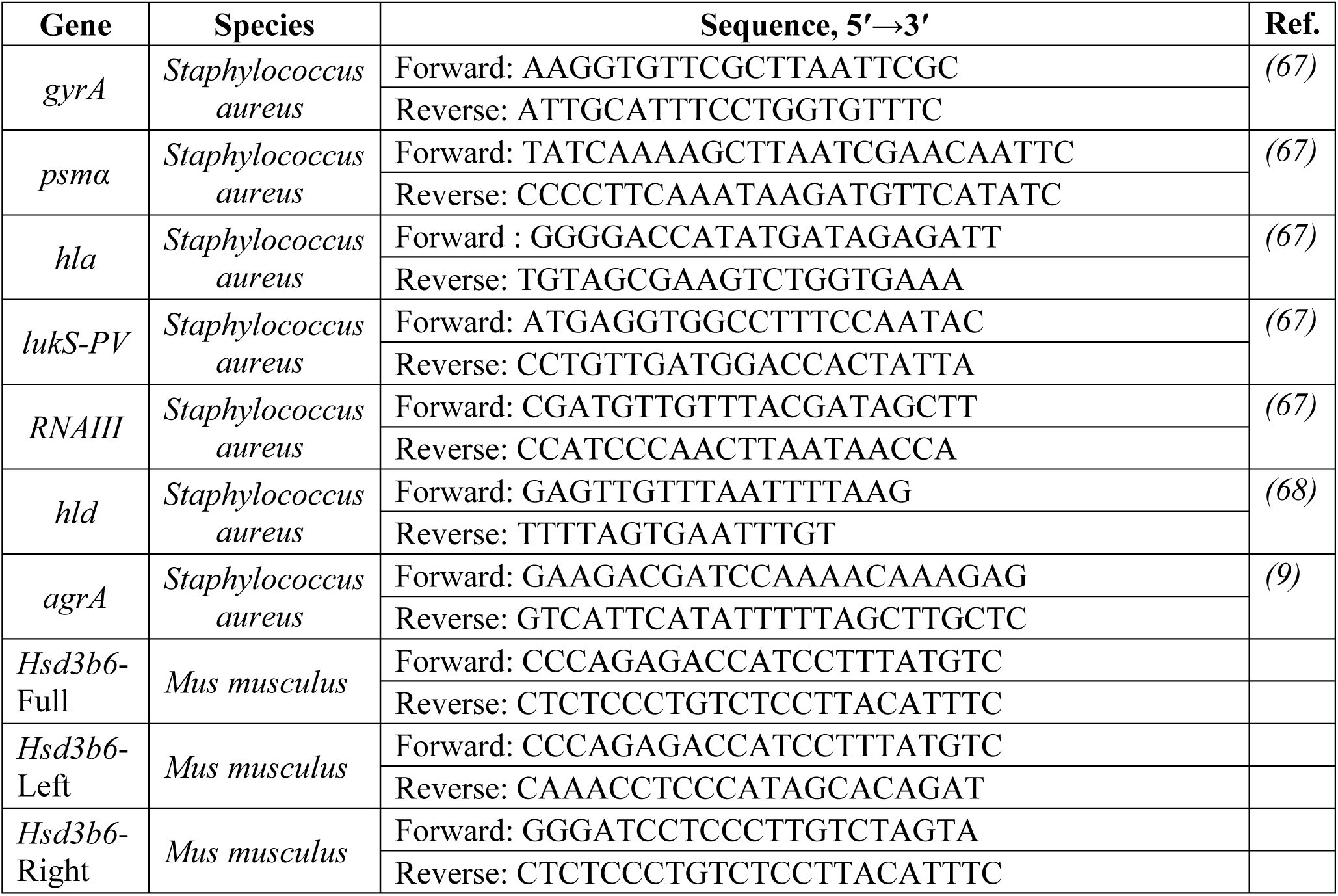
Primers for Quantitative RT-PCR and Gene Sequencing.

